# Excitatory neurons are more disinhibited than inhibitory neurons by chloride dysregulation in the spinal dorsal horn

**DOI:** 10.1101/697417

**Authors:** Kwan Yeop Lee, Stéphanie Ratté, Steven A. Prescott

## Abstract

Neuropathic pain is a debilitating condition caused by the abnormal processing of somatosensory input. Synaptic inhibition in the spinal dorsal horn plays a key role in that processing. Mechanical allodynia – the misperception of light touch as painful – occurs when inhibition is compromised. Disinhibition is due primarily to chloride dysregulation caused by hypofunction of the potassium-chloride co-transporter KCC2. Here we show, in rats, that excitatory neurons are disproportionately affected. This is not because chloride is differentially dysregulated in excitatory and inhibitory neurons, but, rather, because excitatory neurons rely more heavily on inhibition to counterbalance strong excitation. Receptive fields in both cell types have a center-surround organization but disinhibition unmasks more excitatory input to excitatory neurons. Differences in intrinsic excitability also affect how chloride dysregulation affects spiking. These results deepen understanding of how excitation and inhibition are normally balanced in the spinal dorsal horn, and how their imbalance disrupts somatosensory processing.

## INTRODUCTION

Neuropathic pain results from damage to or dysfunction of the nervous system. It affects ∼10% of the population (van Hecke et al., 2014) and is notoriously difficult to treat (Woolf, 2010). Hypersensitivity to tactile stimulation is a troubling feature of such pain. This so-called mechanical allodynia can be acutely reproduced by blocking synaptic inhibition at the spinal level (Miraucourt et al., 2009; Sivilotti and Woolf, 1994; Sorkin and Puig, 1996; Sorkin et al., 1998; Yaksh, 1989). Numerous other studies have shown that synaptic inhibition in the spinal dorsal horn is reduced after nerve injury (for reviews, see Prescott, 2015; Price et al., 2009). Disinhibition can result from reduced activation of GABA_A_ or glycine receptors, or from reduced current flow through activated receptors. The former has several possible causes (Zeilhofer et al., 2012) but the latter stems uniquely from dysregulation of intracellular chloride due to KCC2 hypofunction (Coull et al., 2003). Enhancing KCC2 function reverses injury-induced allodynia (Gagnon et al., 2013; Lavertu et al., 2014; Mapplebeck et al., in press), thus demonstrating that chloride dysregulation contributes significantly to injury-induced disinhibition.

Knowing the molecular mechanism through which disinhibition occurs is important for devising therapeutic interventions, but the effects of disinhibition must also be considered at the circuit level to explain how sensory processing is disrupted (or conversely, to infer how disinhibition arises based on observable/reportable sensory changes that reflect circuit function). Indeed, selective disinhibition of inhibitory or excitatory dorsal horn neurons would likely yield opposite effects. Chloride dysregulation has been shown to occur in dorsal horn neurons with different spiking patterns (Coull et al., 2003), but circuit operation (and sensory processing) depends on how much inhibition each cell type normally experiences and how much is lost under pathological conditions.

Disinhibition is known to unmask low-threshold input to projection neurons that normally respond exclusively to noxious input (Baba et al., 2003; Cheng et al., 2017; Keller et al., 2007; Lavertu et al., 2014; Lu et al., 2013; Miraucourt et al., 2007; Miraucourt et al., 2009; Torsney and MacDermott, 2006). Low-threshold input is conveyed polysynaptically to projection neurons via excitatory interneurons (for reviews, see Peirs and Seal, 2016; Prescott et al., 2014; Takazawa and MacDermott, 2010b). The disproportionate effect of disinhibition on low-threshold input (resulting in allodynia) suggests that excitatory interneurons are disinhibited; greater effects on noxious input (resulting in hyperalgesia) would be expected if disinhibition occurred selectively in projection neurons, i.e. after the convergence of low- and high-threshold inputs. Separate work indicates that excitatory interneurons are necessary for mechanical allodynia (Cheng et al., 2017; Duan et al., 2014; Hu et al., 2006; Malmberg et al., 1997; Wang et al., 2013). Inhibitory neurons also receive inhibitory input (Takazawa and MacDermott, 2010a; Zheng et al., 2010), and could, therefore, also become disinhibited, but whether the resultant increase in GABAergic/ glycinergic transmission enhances inhibition depends on the degree of chloride dysregulation in postsynaptic neurons (Doyon et al., 2011; Prescott et al., 2006). Thus, despite many advances, many questions remain about the balance of excitatory and inhibitory input to different dorsal horn neurons and how this E-I balance becomes pathologically altered.

Beyond “gating” sensory input in the spinal dorsal horn (Melzack and Wall, 1965), synaptic inhibition plays an important role in organizing receptive fields (RFs) (Isaacson and Scanziani, 2011). As originally described in retinal cells by Kuffler (1953), many neurons have an RF with an excitatory center and inhibitory surround. This includes dorsal horn neurons (Hillman and Wall, 1969). A center-surround organization has important implications for sensory processing and for how that processing is disrupted by disinhibition. Notwithstanding other explanations, reduced two-point discrimination in patients with neuropathic pain (Pleger et al., 2006) is consistent with increased RF overlap that could occur if surround inhibition is lost and RFs expand. Animal studies have demonstrated that RFs expand after nerve injury (Behbehani and Dollberg-Stolik, 1994; Cumberbatch et al., 1998; Devor and Wall, 1981; Suzuki et al., 2000; Tabo et al., 1999). This will affect the spatial summation of diffuse (broadly distributed) stimuli – moreso than for punctate (spatially restricted) stimuli – and is liable to affect dynamic mechanical allodynia, which is commonly provoked by clothes touching the skin.

The present study began with an exploration of how chloride dysregulation affects the RFs of superficial dorsal horn neurons. To this end, we blocked spinal KCC2 *in vivo* while monitoring the responses of neurons to tactile stimulation of the hind paw. Beyond RFs expanding, we found that unmasked excitatory input from the original RF surround became the predominant source of excitation. *In vivo* recordings also revealed two distinct cell types: units identified by their adapting spike pattern during tactile stimulation correspond to excitatory neurons whereas non-adapting units correspond to inhibitory neurons. *In vitro* patch clamp recordings showed that chloride regulation did not differ between excitatory and inhibitory neurons, and *in vivo* application of BDNF verified that both cell types experience chloride dysregulation under pathological conditions. That said, adapting and non-adapting cells responded differently to equivalent chloride dysregulation in part because of differences in intrinsic properties and in part because of differences in E-I balance at the circuit level. Most notably, inhibitory neurons were found to receive weak excitation counterbalanced by weak inhibition, whereas excitatory neurons receive strong excitation counterbalanced by strong inhibition. Because excitatory neurons rely more heavily on inhibition, they are disproportionately affected by disinhibition.

## RESULTS

### Disinhibition causes cutaneous receptive fields to expand

For a RF with center-surround organization, disinhibition is expected to expand the RF by unmasking excitatory input originating from the surround (**Fig. 1A**). This will enhance spatial summation, which is critical for diffuse stimuli – like the rubbing of clothes – that typically provoke dynamic allodynia. Compared with the punctate allodynia evoked by stimuli like von Frey hairs, dynamic allodynia is more troubling for patients (Hansson, 2003). Dorsal horn neuron RFs have been reported to change on a short timescale (minutes), consistent with an RF shaped by the balance of excitatory and inhibitory input rather than being “hard wired” (Cook et al., 1987; Dubuisson et al., 1979; Laird and Cervero, 1989; McMahon and Wall, 1984). We therefore set out to determine how the RF size of spinal lamina I-II neurons depends on chloride regulation.

**Figure 1.**
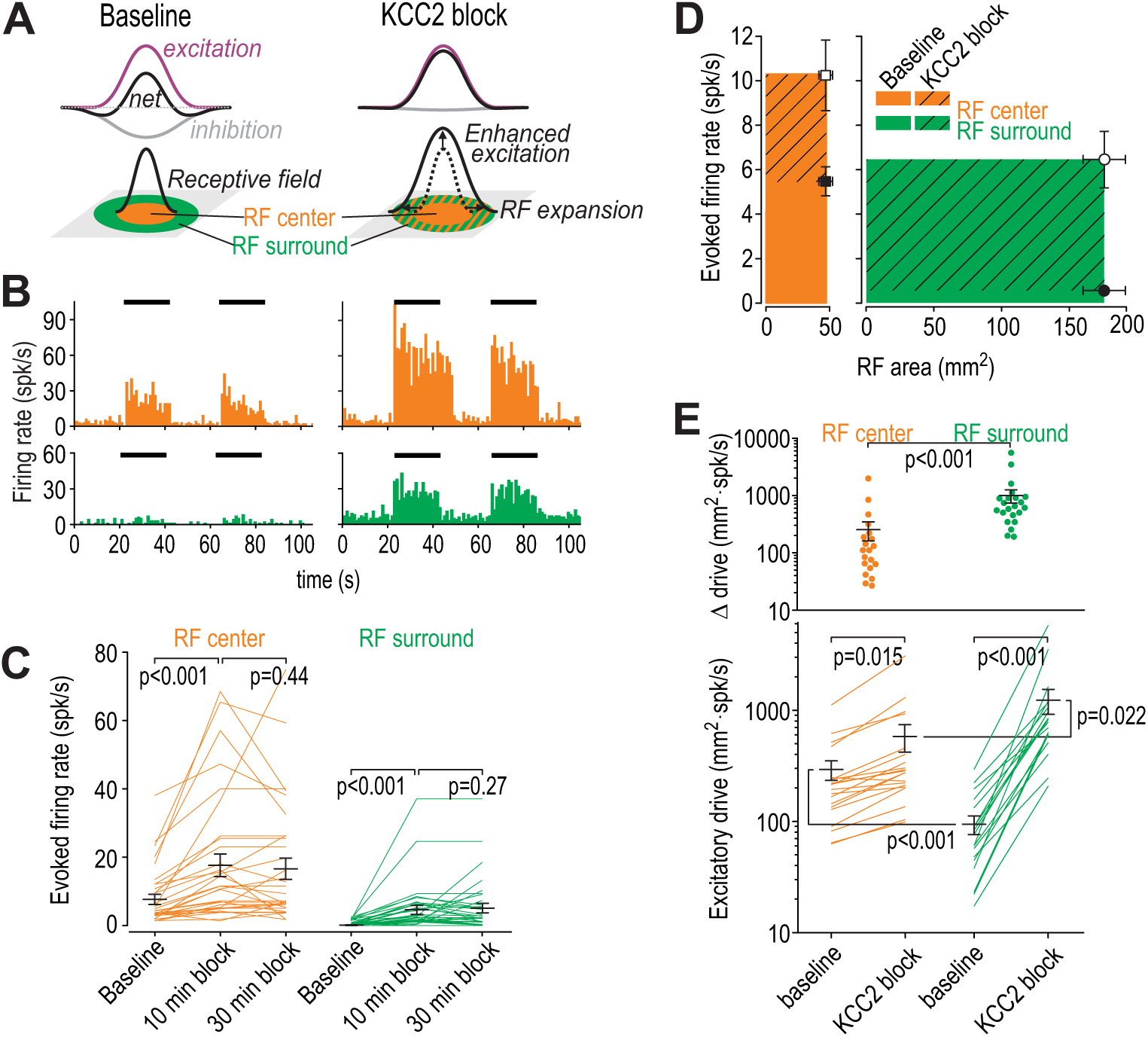
Disinhibition causes receptive fields to expand. **(A)** Cartoon depicts how narrowly tuned excitation combined with broadly tuned inhibition produces a receptive field (RF) with an excitatory center and inhibitory surround. **(B)** Firing rate histograms from a typical neuron responding to brush stimulation (black bars) applied to the RF center (orange) or surround (green) before (left) and after (right) blockade of KCC2 by intrathecal DIOA. **(C)** Summary of spike rates evoked by brush stimulation in the RF center or surround 10 and 30 min after application of 100 µM DIOA (*n* = 10) or 50 µM VU (*n* = 22) to block KCC2; data for DIOA and VU were pooled based on the absence of any significant differences. KCC2 blockade had a significant effect on brush-evoked firing (*F*_2,62_ = 17.78, *p* < 0.001, two-way repeated measures ANOVA). Results of Wilcoxon tests are reported on graphs. Firing rates were stably elevated for at least 20 minutes, thus allowing for other effects of disinhibition (e.g. on RF size) to be measured. **(D)** To quantify the excitatory drive originating from each RF zone, average evoked firing rate (±SEM) was plotted against average surface area (SEM) of the RF zone in which the stimulus was applied. Each data point demarcates a rectangle (colored) whose area is proportional to excitatory drive from that RF zone. **(E)** By calculating the product of RF zone area and firing rate on a cell-by-cell basis (*n* = 22), disinhibition was found to significantly increase total drive (*F*_1,21_ = 139.17, *p* < 0.001; two-way repeated measures ANOVA on log-transformed data) but this depended on the RF zone (interaction, *F*_1,21_= 49.81, *p* < 0.001): the RF surround contributed significantly greater drive than the RF center after KCC2 blockade. Results of Student-Newman-Keuls (SNK) tests are reported on graphs.

To test the effects of KCC2 blockade on RF size, we used brush stimulation to map the cutaneous RF of dorsal horn neurons recorded *in vivo*. The RF measured under normal (baseline) conditions was labeled the RF *center*; by definition, areas outside the RF center evoked negligible spiking (**Fig. 1B** left). After blockade of KCC2 by intrathecal DIOA, stimulation in the RF center evoked larger responses and stimulation outside the RF center evoked vigorous spiking (**Fig. 1B** right). The RF was re-mapped and the additional area was labeled the RF *surround*. KCC2 blockade significantly increased firing evoked by stimulation in the RF center or surround (*Z* = 4.79, *p* < 0.001 and *Z* = 4.72, *p* < 0.001 respectively; Wilcoxon tests) (**Fig. 1C**). Firing rate histograms in Figure 1B include all spikes but the spontaneous firing rate preceding each stimulus was subtracted from firing during stimulation to focus on evoked firing in Figure 1C-E.

Next, we compared the enhanced excitatory drive from the RF center with the unmasked excitatory drive from the RF surround by plotting the average area for each RF zone against the average firing rate evoked by stimulation in that zone. This defines a rectangle whose area is proportional to the total excitatory drive contributed by that RF zone (**Fig. 1D**). By multiplying the area of each RF zone with the corresponding firing rate for each unit (i.e. calculating the area of the aforementioned rectangle on a cell-by-cell basis), we found that KCC2 blockade significantly increased drive from both the RF center and surround (*Q*_21_ = 3.60, *p* = 0.015 and *Q*_21_ = 18.54, *p* < 0.001, respectively; post-hoc Student-Newman-Keuls [SNK] tests). Excitatory drive from the RF surround was (by definition) significantly less than from the RF center under baseline conditions (*Q*_21_ = 5.91, *p* < 0.001), but the inverse was true after disinhibition (*Q*_21_ = 3.42, *p* = 0.022) (**Fig. 1E**). Hence, after disinhibition caused by KCC2 blockade, neurons were driven more strongly by unmasked excitatory input from the RF surround than by enhanced input from the RF center.

### Units recorded in vivo comprise two distinct groups

Over the course of these experiments we noticed that some units had a monophasic spike waveform whereas others had a biphasic waveform (**Fig. 2A**). The former exhibited spike rate adaptation during sustained tactile stimulation with a von Frey (VF) filament whereas the latter did not (**Fig. 2B**). The adaptation time constant measured from exponential-rise-to-maximum fits of the cumulative probability distribution of spike times from each unit differed significantly between units classified by their spike waveform (*T*_23_ = 8.58, *p* < 0.001; unpaired *t*-test) (**Fig. 2B** inset). The difference in adaptation argues that units with distinct spike waveforms represent distinct cell types rather than, for example, somatic and axonal recording from the same population of neurons. The two cell types are henceforth referred to as *adapting* and *non-adapting*. In subsequent experiments, we found that approximately 80% of units responsive to light tactile stimulation could be readily classified by their adaptation pattern; unclassified units were excluded from subsequent analysis.

**Figure 2.**
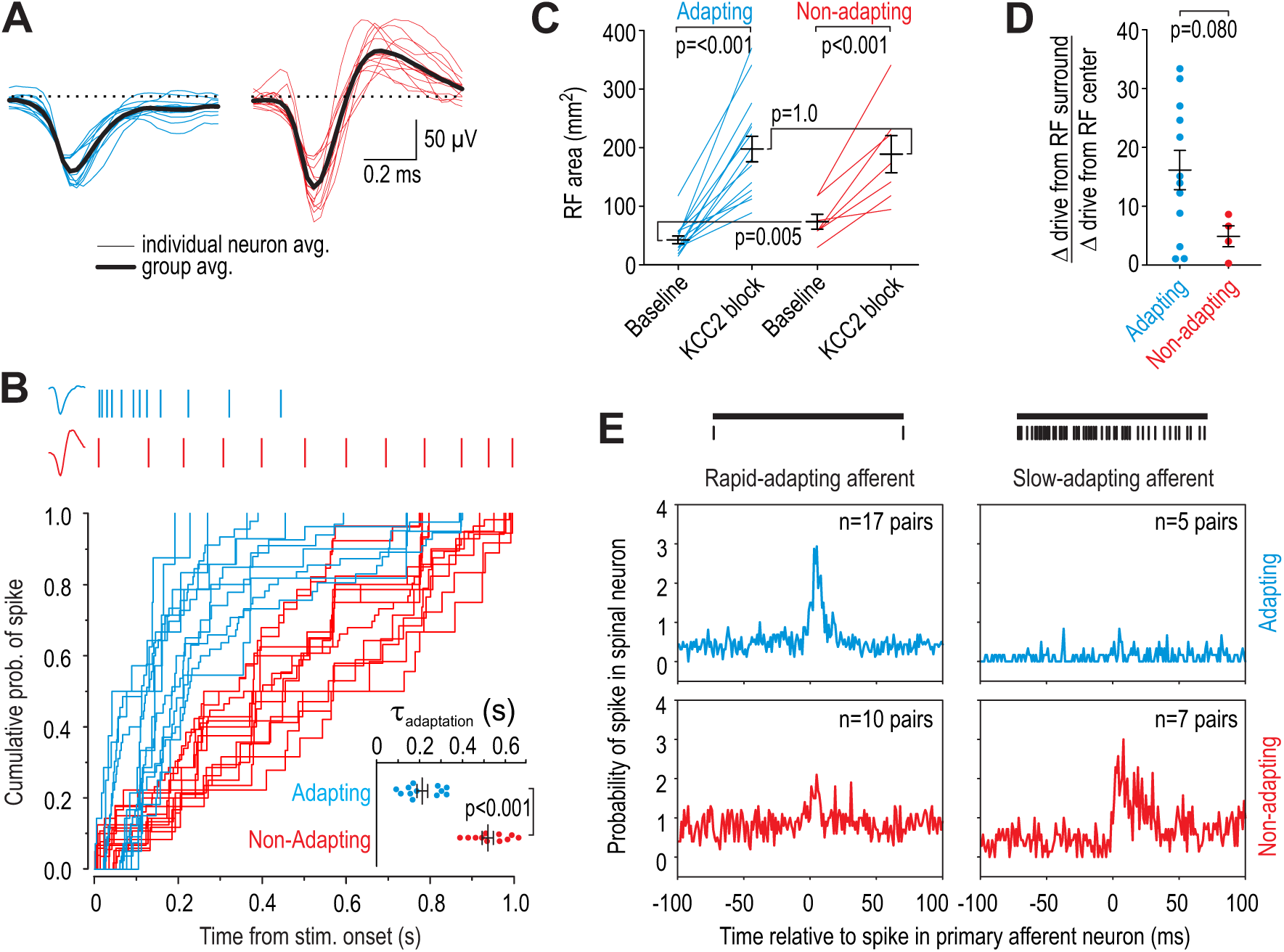
Units recorded *in vivo* can be divided into two groups. **(A)** Spike waveforms were separated according to their monophasic (blue) or biphasic (red) shape. **(B)** Units subdivided by spike waveform also differed in their adaptation pattern, as illustrated by sample rasters showing responses to 1 s-long 10 g von Frey stimulation (top). The cumulative probability of spiking was fit for each unit with the curve *y* = 1/(1-e^-t/*τ*^). The adaptation time constant *τ* differed significantly between groups (*T*_23_ = 8.58, *p* < 0.001; unpaired *t*-test). Units are henceforth referred to as adapting (blue) or non-adapting (red). **(C)** Adapting units had a significantly smaller RF than non-adapting units at baseline (*Q*_21_ = 17.5, *p* = 0.013; Mann-Whitney test) but there was no difference in RF size after KCC2 blockade (*Q*_21_ = 49.0, *p* = 0.83). RF size was significantly affected by KCC2 blockade in both type of units (*F*_1,20_ = 66.10, *p* < 0.001; two-way repeated measures ANOVA). Results of SNK tests are reported on graphs. **(D)** KCC2 blockade caused a larger, though not quite significant, increase in drive from the RC center relative to the RF surround in adapting units compared with non-adapting units (*T*_14_ = 1.89, *p* = 0.080; unpaired *t*-test). **(E)** Cross-correlograms were calculated from all pairs of simultaneously recorded primary afferent neurons and spinal neurons with overlapping RFs. The number of pairs is reported on each panel. Primary afferents were classified as rapid-adapting (RA, left) or slow-adapting (SA, right); rasters show sample responses to 1 s-long 10 g von Frey stimulation (top). Adapting units displayed an increased probability of spiking after RA spikes (top left) but not after SA spikes (top right) whereas non-adapting units displayed a large increase in spike probability after SA spikes (bottom right) and a modest increase after RA spikes (bottom left). In other words, adapting units receive input exclusively from RA afferents whereas non-adapting units receive input preferentially from SA afferents.

Analysis of additional response properties support the classification based on adaptation pattern. The RFs of adapting units were significantly smaller than RFs of non-adapting units at baseline (*U* = 17.5, *p* = 0.013; Mann-Whitney test) (**Fig. 2C**). KCC2 blockade enlarged RFs in both adapting units (*Q*_14_ = 11.69, *p* < 0.001; SNK test) and non-adapting units (*Q*_6_ = 5.94, *p* < 0.001). Of the units analyzed in Figure 1E, 12 were subsequently identified as adapting and 4 as non-adapting; the disinhibition-mediated increase in drive from the RF surround relative to increased drive from the RF center tended to be greater in adapting units, though the difference did not reach significance (*T*_14_ = 1.89, *p* = 0.080; unpaired *t*-test) (**Fig. 2D**).

To assess functional connectivity between primary afferent neurons and each type of spinal unit, we recorded simultaneously from the dorsal root ganglion and spinal cord. Mechanosensitive afferents were subdivided into rapid-adapting and slow-adapting (Johansson and Vallbo, 1979). We calculated cross-correlograms by measuring the probability of a spinal unit firing at different times relative to each spike in a simultaneously recorded primary afferent with overlapping RF. According to this analysis, adapting units received tactile input exclusively from rapid-adapting afferents whereas non-adapting units received tactile input preferentially from slow-adapting afferents (**Fig. 2E**). Differences in adaptation during evoked firing are likely due, at least in part, to differences in primary afferent input, but differences in spontaneous spiking (see Fig. 4C) suggest that adapting and non-adapting units also differ in their intrinsic excitability (see below).

**Figure 3.**
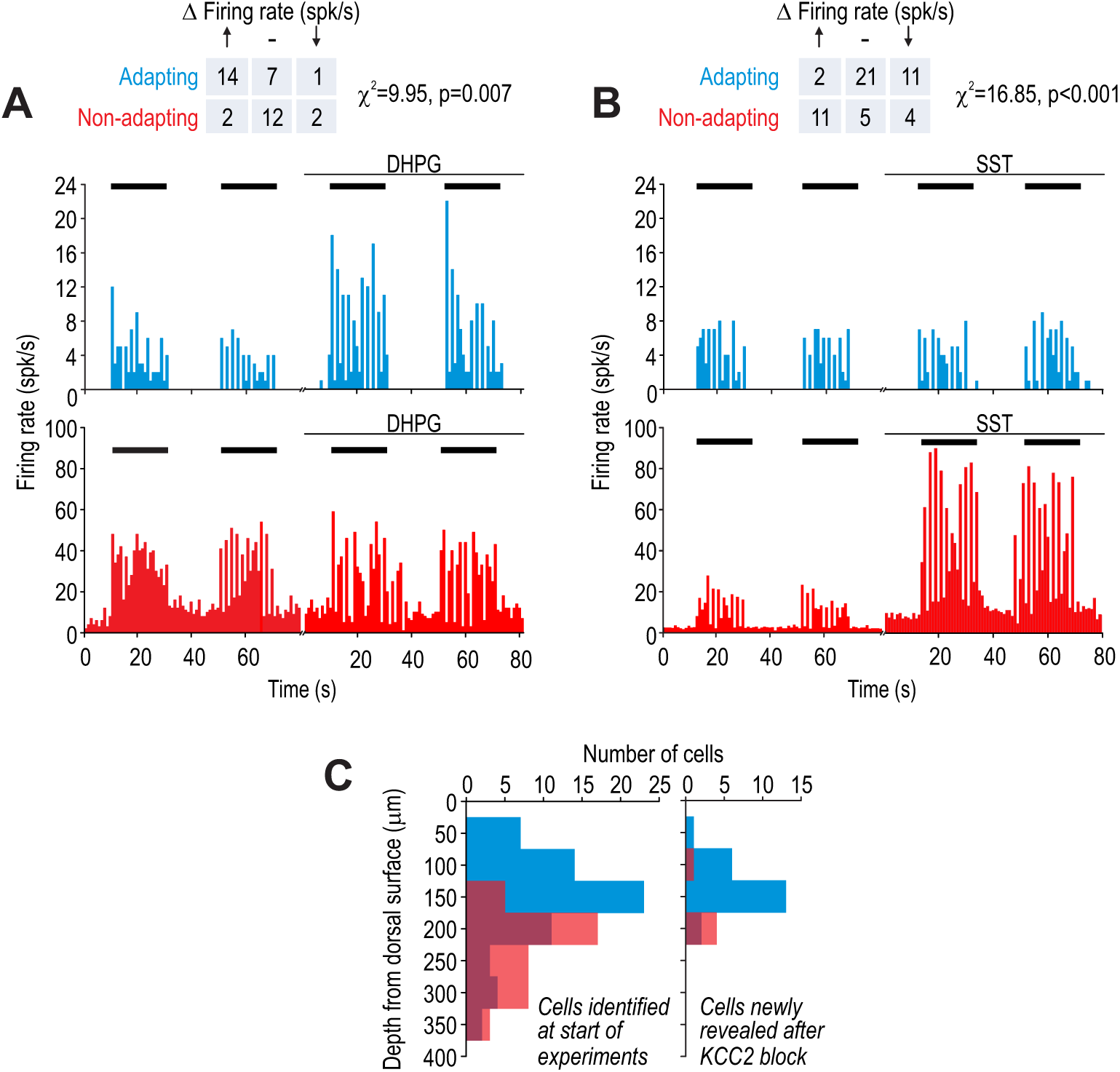
Adapting and non-adapting units correspond to excitatory and inhibitory neurons, respectively. Sample firing rate histograms from a typical adapting unit (blue) and non-adapting unit (red) during brush stimulation (bars) before and after intrathecal application of 200 µM 3,5-dihydroxyphenylglycine (DHPG). Adapting units were disproportionately affected (χ^2^ = 9.95, *p* = 0.007), consistent with them being excitatory neurons, which selectively express type 5 metabotropic glutamate receptors. **(B)** Sample firing rate histograms before and after intrathecal application of 50 µM somatostatin (SST). Non-adapting units were disproportionately affected (χ^2^ = 16.85, *p* = 0.001), consistent with them being inhibitory neurons, which selectively express the sst_2a_ receptor. Some adapting units exhibited reduced spiking (see Results). Spontaneous firing was increased by SST selectively in non-adapting units. **(C)** Distribution of recording depths for units responsive to light touch at baseline (left) and for units that became responsive after KCC2 blockade (right).

**Figure 4.**
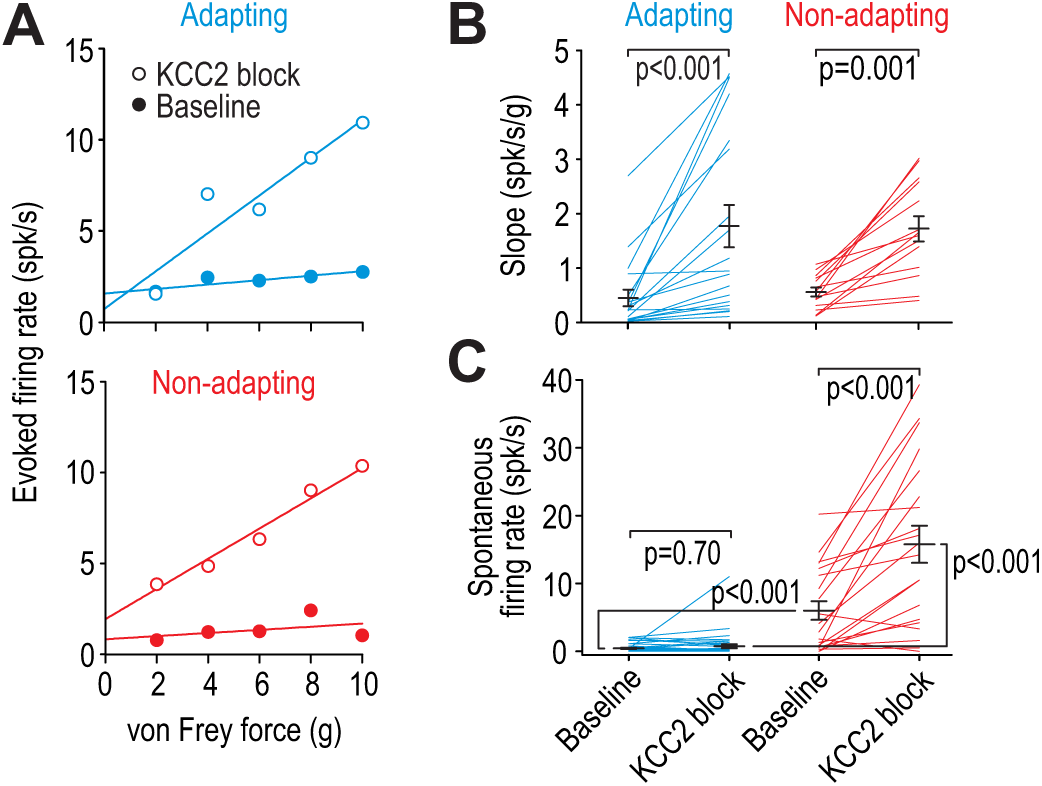
Adapting and non-adapting units are differentially affected by KCC2 blockade. **(A)** Input-output (*i-o*) curves show firing rate evoked by increasing force applied by von Frey hairs to the RF surround in a typical adapting unit (blue) and non-adapting unit (red) before and after blockade of KCC2. **(B)** KCC2 blockade had a significant effect on slope (*F*_1,31_ = 35.68, *p* < 0.001; two-way repeated measures ANOVA). Results of Wilcoxon tests are reported on graphs. **(C)** KCC2 blockade also had a significant effect on spontaneous firing (*F*_1,57_ = 58.60, *p* < 0.001; two-way repeated measures ANOVA) but that effect differed between cell types (*F*_1,57_ = 33.79, *p* < 0.001): non-adapting units experienced a significant increase in spontaneous firing (*Z*_19_ = 3.58, *p* < 0.001; Wilcoxon test) whereas adapting units did not (*Z*_38_ = 0.39, *p* = 0.70). Spontaneous firing differed significantly between cell types at baseline (*U*_58_ = 116.5, *p* < 0.001; Mann-Whitney test) and after KCC2 blockade (*U*_58_ = 55.5, *p* < 0.001).

### Adapting and non-adapting units correspond to excitatory and inhibitory neurons, respectively

Next, we asked whether adapting and non-adapting units represent different cell types within the dorsal horn circuit. Since projection neurons are absent in lamina II and represent only ∼5% of neurons in lumbar segments of lamina I in rats (Spike et al., 2003), the units we recorded were most likely local interneurons, of which approximately 2/3 are excitatory and 1/3 are inhibitory (Abraira et al., 2017; Polgar et al., 2003; Prescott and Ratté, 2012; Todd, 2010, 2017). Without transgenic rats to genetically identify neuron types, and rather than switch to mice in which certain experiments (e.g. Figs. 2E or 8B) would be prohibitively difficult, we undertook a series of tests designed to link adaptation pattern to pharmacologically testable markers and other distinctive properties.

First, we intrathecally applied agonists of receptors expressed selectively by excitatory or inhibitory dorsal horn neurons to test for differential effects in adapting and non-adapting units. Metabotropic glutamate receptor 5 is expressed almost exclusively on excitatory neurons and its agonist 3,5-dihydroxyphenylglycine (DHPG) has been shown to selectively modulate the excitability of excitatory neurons (Hu and Gereau, 2011). Comparing brush-evoked firing before and after intrathecal DHPG revealed that adapting cells were preferentially affected (χ^2^ = 9.95, *p* = 0.007) (**Fig. 3A**), consistent with adapting cells being excitatory neurons. On the other hand, the somatostatin (SST) receptor sst_2a_ is expressed selectively on inhibitory neurons (Todd et al., 1998). Intrathecal application of SST preferentially affected non-adapting cells (χ^2^ = 16.85, *p* < 0.001) (**Fig. 3B**), consistent with non-adapting cells being inhibitory neurons. Unlike most non-adapting units whose evoked firing was increased by SST, evoked firing was reduced in a minority of adapting units presumably via an indirect circuit-level effect, namely, increased inhibition mediated by sst_2a_-expressing neurons. Non-adapting units also exhibited an increase in spontaneous firing (see Fig. 3B), which must be a direct effect of SST on those units given the absence of SST-induced spontaneous firing in any other units.

**Figure 3C** summarizes the depth of each unit from the dorsal surface of the spinal cord. The distributions of adapting and non-adapting units were significantly different (*D* = 0.56, *p* < 0.001; Kolmogorov-Smirnov test) with an average depth (± SD) of 156 ± 72 µm and 234 ± 57 µm, respectively (**Fig. 3C** left), which is consistent with the increasing proportion of inhibitory neurons between lamina I to III (Polgar et al., 2003). The relative numbers of adapting and non-adapting units (*n* = 64 and 41, respectively) did not differ significantly (χ^2^ = 0.74, *p* = 0.39) from the overall 2:1 ratio of excitatory to inhibitory neurons in the superficial dorsal horn (see above), though one must consider that we recorded selectively from neurons responsive to innocuous tactile stimulation. We identified an additional 27 units that responded to tactile stimulation only after KCC2 blockade (**Fig. 3C** right). These units presumably receive low-threshold input via polysynaptic pathways normally blocked by inhibition. Of these units, 22 were adapting and 5 were non-adapting, consistent with the emerging picture that polysynaptic pathways comprising excitatory interneurons can relay low-threshold inputs to superficial projection neurons (see Introduction).

### Spiking in adapting and non-adapting units is differentially affected by KCC2 blockade

To explore whether putative excitatory and inhibitory neurons are differentially affected by chloride dysregulation, we measured evoked firing before and after KCC2 blockade. **Figure 4A** shows sample input-output (*i-o*) curves for representative adapting and non-adapting units activated by von Frey stimulation of their RF surround. KCC2 blockade significantly increased *i-o* curve slope in adapting and non-adapting units (*Z* = 3.82, *p* < 0.001 and *Z* = 3.30, *p* < 0.001, respectively; Wilcoxon tests) (**Fig. 4B**). The *y*-intercept was not analyzed since the spontaneous firing rate was subtracted to isolate the evoked response, but separate analysis showed that spontaneous firing was significantly greater in non-adapting units than in adapting units at baseline (*U* = 116.5, *p* < 0.001; Mann-Whitney test), and that KCC2 blockade significantly increased spontaneous firing in non-adapting units (*Z* = 3.58, *p* < 0.001; Wilcoxon test) but not in adapting units (*Z* = 0.39, *p* = 0.70) (**Fig. 4C**). Recall that spontaneous firing in non-adapting units was also increased by SST (see Fig. 3B), yet other manipulations like glycine receptor blockade did not affect spontaneous firing in non-adapting units (*T*_9_ = 1.654, *p* = 0.13; paired *t*-test) (**Fig. S4**). The latter observation argues that spontaneous firing is not a consequence of circuit-level disinhibition, but is instead most likely due to the depolarization caused by manipulations like KCC2 blockade. Indeed, inhibitory neurons are under strong tonic GABAergic or glycinergic inhibition (Gradwell et al., 2017; Takazawa and MacDermott, 2010a) whose disruption can cause depolarization, and tonic-spiking inhibitory neurons respond to subtle depolarization with sustained spiking because of their voltage-gated channels (Prescott and De Koninck, 2005; Ratté et al., 2015). To summarize, KCC2 blockade affected touch-evoked spiking in both adapting and non-adapting units but the selective modulation of spontaneous spiking in the latter likely reflects the intrinsic properties of that cell type.

### Excitatory and inhibitory neurons are differentially affected by equivalent chloride dysregulation

Differential effects of KCC2 blockade on spiking in adapting and non-adapting units could be due to differences in chloride regulation (KCC2 expression) and/or differences in intrinsic excitability that influence how chloride dysregulation affects spiking. To test the former, we conducted whole-cell patch clamp recordings in a spinal slice preparation (see Methods). Specifically, we identified neurons as excitatory or inhibitory based on their spiking pattern recorded in current clamp (see below) and, after loading the neuron with high chloride, we measured chloride reversal potential (*E*_Cl_) before and after blocking KCC2 based on voltage clamp responses to puffed GABA (**Fig 5A**). Changes in *E*_Cl_ measured under a defined chloride load is a more sensitive measure of chloride extrusion capacity than perforated patch measurements of *E*_Cl_ in low chloride-load conditions (Doyon et al., 2016). Results revealed no difference in chloride extrusion capacity between excitatory and inhibitory neurons (*T*_12_ = 0.030, *p* = 0.98, unpaired *t*-test) (**Fig. 5B**), thus excluding differences in chloride regulation as the basis for the differentially modulated spiking reported in Figure 4C.

**Figure 5.**
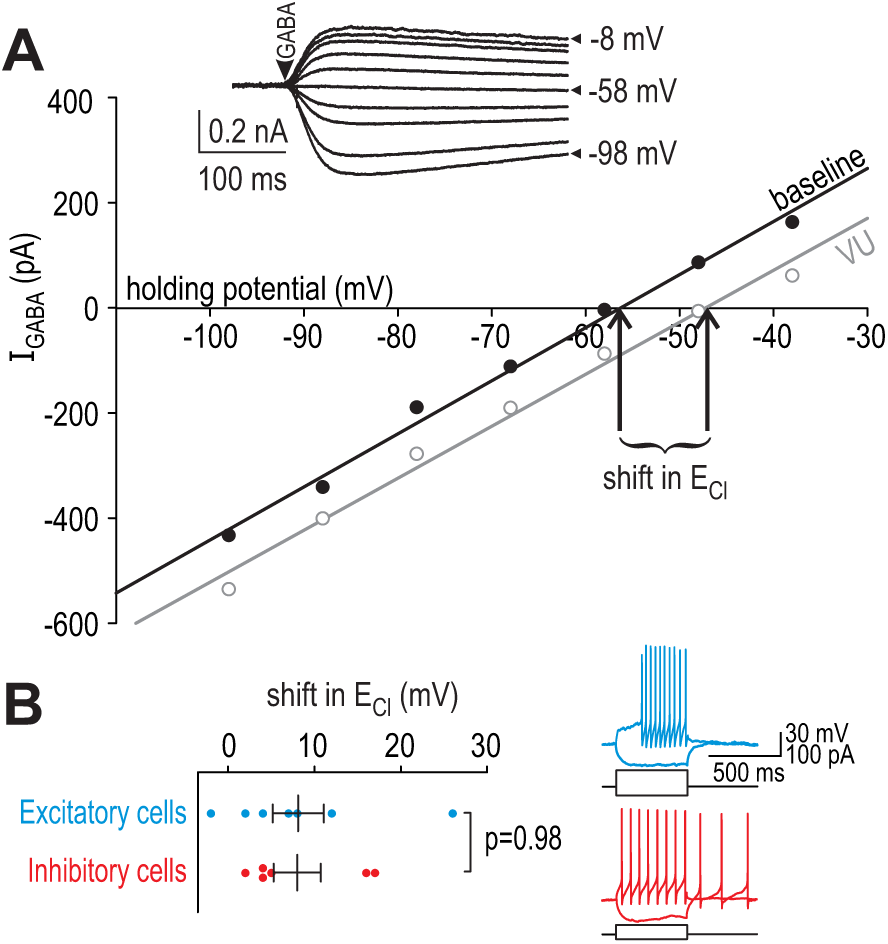
Excitatory and inhibitory neurons have equivalent chloride extrusion capacity. **(A)** Responses to 5-10 ms-long puffs of 1 mM GABA (arrowhead) were measured in voltage clamp at different holding potentials (inset) in neurons patched with a high-chloride (25 mM) pipette solution. Chloride reversal potential *E*_Cl_, which corresponds to where the *I-V* curve intersects the *x*-axis, was measured before (black) and after (gray) blockade of KCC2 with 15 µM bath-applied VU. The shift in *E*_Cl_ reflects chloride extrusion capacity. **(B)** Neurons identified as excitatory or inhibitory based on spiking pattern (inset) displayed no difference in chloride extrusion capacity (*T*_12_= 0.030, *p* = 0.98; unpaired *T*-test).

For all *in vitro* experiments, excitatory and inhibitory neurons were identified by their spiking pattern during sustained somatic current injection (see traces on **Fig. 5B**). The A-type potassium current, which is associated with delayed- and single-spiking (Balachandar and Prescott, 2018), is selectively expressed in the spinal dorsal horn by excitatory neurons (Hu et al., 2006) whereas tonic-spiking is observed almost exclusively in inhibitory neurons (Yasaka et al., 2010) (for reviews, see Prescott and Ratté, 2012; Todd, 2010, 2017). This is consistent with excitatory neurons genetically identified by somatostatin expression (most of which exhibit delayed-, single-, or burst-spiking) and inhibitory neurons genetically identified by dynorphin expression (most of which exhibit tonic- or burst-spiking) (Duan et al., 2014). Thus, although some spiking patterns (e.g. bursting) are not predictive of cell type, other patterns are (see also Hughes et al., 2012). Accordingly, we tested single- or delayed-spiking excitatory neurons and tonic-spiking inhibitory neurons; neurons with other spiking patterns or whose classification was unclear were excluded. Given these inclusion criteria, data in Figure 5 do not rule out differences in chloride extrusion within certain subpopulations of neurons, but they do rule out a systematic difference between the most common excitatory and inhibitory neurons.

To test whether differences in intrinsic excitability influence how chloride dysregulation affects spiking, we used dynamic clamp to simulate equivalent virtual disinhibition in different cell types. All neurons received virtual excitatory and inhibitory synaptic inputs modeled as Ornstein-Uhlenbeck processes (**Fig. 6A**). Firing rate was measured for different average excitatory conductance (*g*_exc0_). The average inhibitory conductance (*g*_inh0_) was co-varied with *g*_exc0_ according to α, where α = ½ represents stimulation inside the RF whereas α = 2 represents stimulation in the RF surround (**Fig. 6B**). This parameterization assumes a center-surround organization formed by the combination of narrowly tuned excitatory input and broadly tuned inhibitory input (see Introduction). Chloride dysregulation was modeled as a shift in the anion reversal potential (*E*_inh_) from −70 mV to −45 mV (Coull et al., 2003).

**Figure 6.**
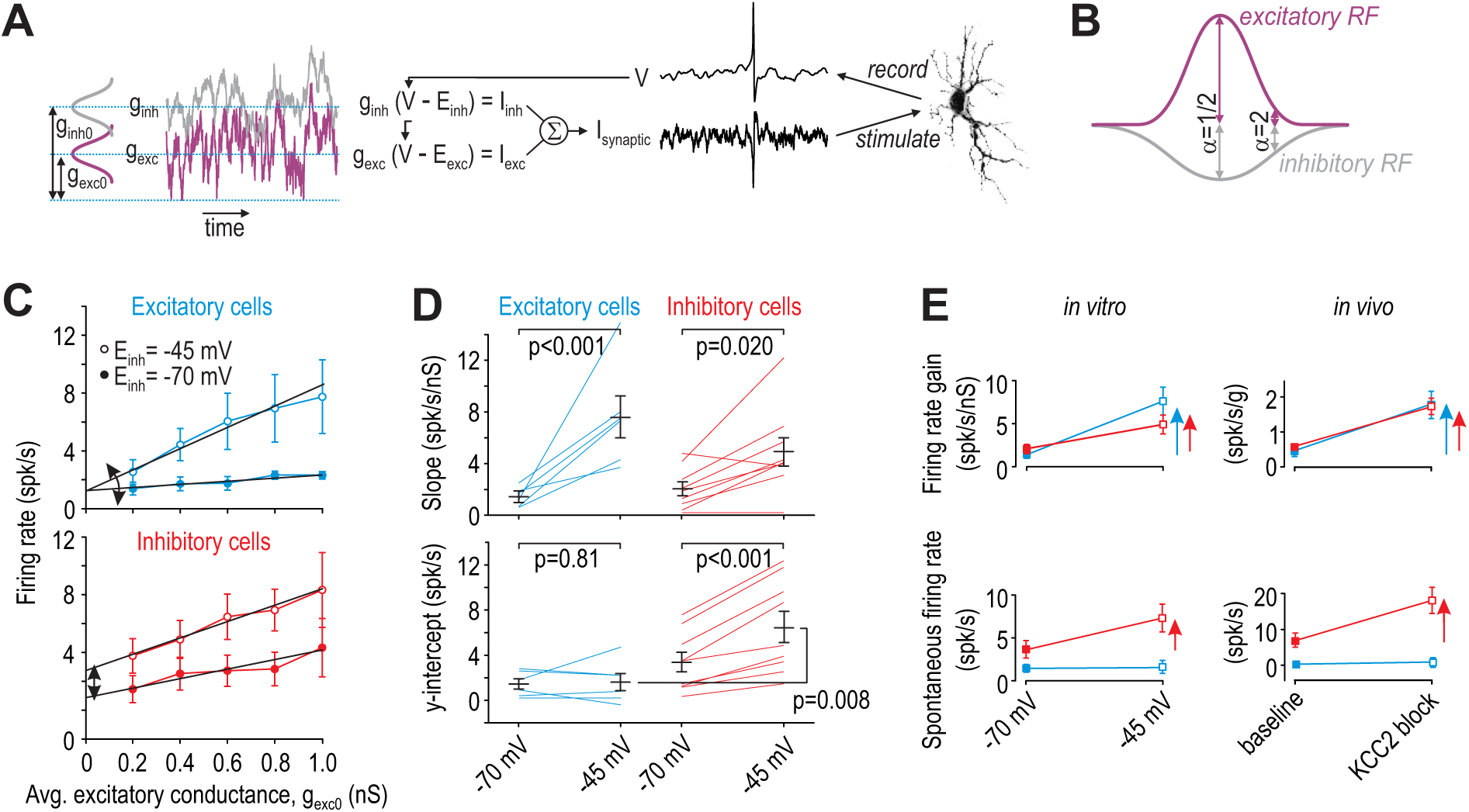
Excitatory and inhibitory neurons are differentially affected by virtual disinhibition. **(A)** Cartoon depicts dynamic clamp experiments: noisy excitatory and inhibitory conductances were generated through separate Ornstein-Uhlenbeck processes, converted to currents based on recorded voltage, and applied to the neuron. Average excitatory conductance, *g*_exc0_, was systematically varied to plot input-output (*i-o*) curves. **(B)** Average inhibitory conductance, *g*_inh0_, was co-varied with *g*_exc0_ according to α = *g*_inh0_/*g*_exc0_, where α = 0.5 for virtual stimulation in the RF center and α = 2 for virtual stimulation in the RF surround. **(C)** Average *i-o* curves (±SEM) for excitatory cells (blue, *n* = 6) and inhibitory cells (red, *n* = 8) for = 2; see Fig. S6 for results with = 0.5. Each neuron was tested with normal inhibition (*E*_inh_ = −70 mV) and again after virtual disinhibition (*E*_inh_ = −45 mV). Disinhibition affected the *i-o* curve slope of both cell types but shifted the *i-o* curve only in inhibitory cells. **(D)** *E*_inh_ had a significant effect on slope (*F*_1,13_ = 28.12, *p* < 0.001, two-way repeated measures ANOVA). *E*_inh_ also had a significant effect on the *y*-intercept (*F*_1,13_ = 14.48, *p* = 0.002) but only in inhibitory neurons (interaction, *F*_1,13_ = 11.75, *p* = 0.004). Results of SNK tests are reported on graphs. **(E)** Summary of disinhibitory effects *in vitro* (data from panels C and D) for comparison with disinhibitory effects *in vivo* (data from Figure 4). Data are summarized as mean ± SEM. Each colored arrow indicates a significant effect of disinhibition (*p* < 0.05). Disinhibition significantly increased firing rate gain in all neuron types but increased spontaneous firing selectively in inhibitory neurons and non-adapting units.

**Figure 6C** shows the average *i-o* curves for excitatory neurons (top) and inhibitory neurons (bottom) tested with α = 2. Both types of neurons responded to virtual chloride dysregulation with an increase in firing rate gain (slope) but only inhibitory neurons exhibited a pronounced shift in their average *i-o* curve. To quantify these changes, we fitted data from each neuron individually and compared the slope and *y*-intercept between cell types. Disinhibition significantly increased the slope in both excitatory and inhibitory neurons (*Q*_5_ = 6.62, *p* < 0.001 and *Q*_8_ = 3.75, *p* = 0.020, respectively; SNK tests) (**Fig. 6D** top), but increased the *y*-intercept only in inhibitory neurons (*Q*_8_ = 8.09, *p* < 0.001 vs *Q*_5_ = 0.35, *p* = 0.81 in excitatory neurons) (**Fig. 6D** bottom). Similar trends were observed for α = ½ (**Fig. S6**) though effects of disinhibition were weaker, consistent with weaker inhibition relative to excitation in the RF center. Indeed, firing rate gain under normal conditions (*E*_inh_ = −70 mV) was significantly less for α = 2 than for α = ½ (*Q*_5_ = 8.67, *p* < 0.001 and *Q*_5_ = 7.45, *p* < 0.001 for excitatory and inhibitory neurons, respectively; SNK tests), consistent with past work on inhibition-mediated gain control (Prescott and De Koninck, 2003; Silver, 2010).

**Figure 6E** summarizes the effects of virtual disinhibition *in vitro* (left panels) for comparison with effects of KCC2 blockade *in vivo* (right panels). Whereas the firing rate gain of all cell types was significantly increased by disinhibition, only inhibitory neurons (*in vitro*) and non-adapting units (*in vivo*) exhibited a significant change in spontaneous spiking. The concordance between *in vitro* and *in vivo* data supports the interpretation of data in Figure 3, namely that adapting units represent excitatory neurons whereas non-adapting units represent inhibitory neurons.

### BDNF induces chloride dysregulation in both cell types

Although excitatory and inhibitory neurons exhibit equivalent chloride extrusion capacity according to Figure 5, those results do not address whether KCC2 is equivalently downregulated in each cell type after nerve injury. To address this issue, we tested whether adapting and non-adapting units are comparably affected by brain-derived neurotrophic factor (BDNF), which plays a key role in downregulating KCC2 after nerve injury (Coull et al., 2005). Notwithstanding sex differences in neuroimmune signaling (Sorge et al., 2015), we recently showed that BNDF is sufficient to downregulate KCC2 and cause allodynia in both male and female rodents (Mapplebeck et al., in press). Rather than compare nerve-injured and control animals, in which other changes induced by nerve injury might compromise our classification scheme, we compared responses for each unit before and after intrathecal BDNF, and re-tested again after applying acetazolamide (ACTZ), which selectively reverses effects of chloride dysregulation by reducing bicarbonate efflux through GABA_A_/glycine receptors (Lee and Prescott, 2015). Reversal of BDNF effects by ACTZ serves as a control for off-target (non-KCC2-mediated) effects of BDNF.

Intrathecal BDNF significantly increased firing evoked by brush stimulation applied to the RF center of adapting or non-adapting units (*Z* = 3.10, *p* < 0.001 for both cell types; Wilcoxon tests) (**Fig. 7A** left). Comparable effects were observed for stimuli applied to the RF surround (*Z* ≥ 2.93, *p* < 0.001) (**Fig. 7A** right). In all cases, BDNF-mediated enhancement of evoked firing was significantly reversed by subsequent application of ACTZ (*Z* ≥ 2.85, *p* ≤ 0.002). Moreover, BDNF failed to induce spontaneous spiking in adapting units (*Z* = 0.51 *p* = 0.64) but significantly increased spontaneous firing in non-adapting units (*Z* = 2.52, *p* = 0.008) (**Fig. 7B**). This differential change in spontaneous firing is consistent with the effect of KCC2 blockade (Fig. 4C), and is unlike the effect of glycine receptor blockade (Fig. S4). The BDNF-mediated enhancement of spontaneous firing was significantly reversed by ACTZ (*Z* = 2.52, *p* = 0.008). These results demonstrate that BDNF causes comparable disinhibition in both cell types, but, like for direct blockade of KCC2, spontaneous firing was increased selectively in non-adapting units.

**Figure 7.**
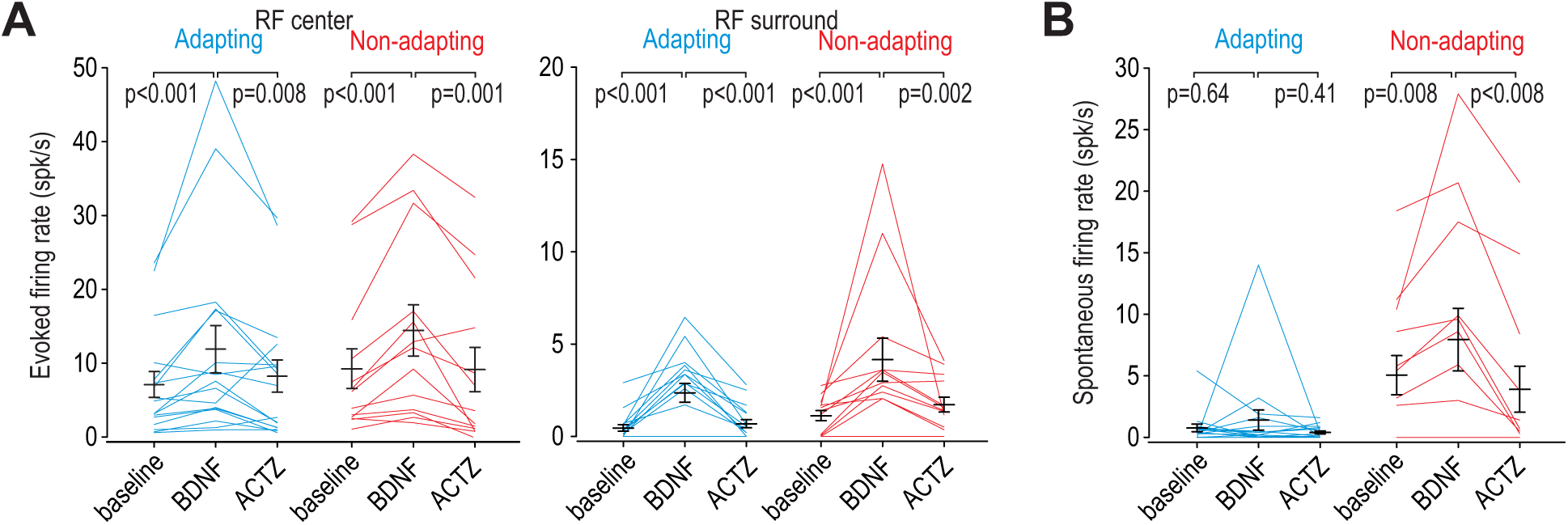
BDNF causes disinhibition in both adapting and non-adapting units. **(A)** Intrathecal BDNF (70 µg) and ACTZ (10 mM) had a significant effect on the firing evoked by brush stimulation of the RF center (*F*_2,56_ = 15.41, *p* < 0.001; two-way repeated measures ANOVA) or surround (*F*_2,56_ = 17.92, *p* < 0.001). BDNF significantly increased evoked firing and subsequent application of ACTZ significantly reduced firing in adapting (blue) and non-adapting (red) units. Results of SNK tests are reported on graphs. **(B)** BDNF and ACTZ also had a significant effect on spontaneous firing (*F*_2,56_ = 6.76, *p* = 0.002; two-way repeated measures ANOVA) but that effect differed between cell types (interaction, *F*_1,56_ = 8.10, *p* = 0.008). Only non-adapting units responded to BDNF with significantly increased spontaneous firing, which was entirely reversed by ACTZ. Results of SNK tests are reported on graphs.

### Surround inhibition is greater, and disininhibition more consequential, for adapting units

In a final set of experiments, we tested how expansion of the RF affects spatial summation of tactile input. To this end, we compared responses to co-stimulation in the RF center *and* surround with responses to stimulation in the RF center alone. Stimulation of the RF surround is predicted to reduce evoked firing under baseline conditions (ratio <1) and to increase it after KCC2 blockade (ratio >1) (**Fig. 8A**). As predicted, stimulation in the RF surround significantly reduced the response to stimulation in the RF center (*T*_22_ = −9.26, *p* < 0.001 and *T*_11_ = −2.90, *p* = 0.014 for adapting and non-adapting units, respectively; one sample *t-*tests on log-transformed data) (**Fig. 8B**). Adapting units experienced a significantly larger reduction than non-adapting units (*Q*_33_ = 3.56, *p* = 0.014; SNK test). After KCC2 blockade, the effect of stimulation in the RF surround was inverted: Stimulation in the RF surround significantly enhanced the response to stimulation in the RF center (*T*_22_ = 6.94, *p* < 0.001 and *T*_11_ = −4.58, *p* < 0.001 for adapting and non-adapting units, respectively). Moreover, adapting units experienced a significantly larger enhancement that non-adapting units (*Q*_33_ = 3.78, *p* = 0.010; SNK test). This pattern reveals that adapting units receive strong excitatory input balanced by strong inhibitory input, whereas non-adapting units receive weaker excitatory input balanced by weaker inhibitory input (**Fig. 8C**). Because of their heavier reliance on inhibition to counterbalance strong excitation, adapting units experience greater disinhibition than non-adapting units.

**Figure 8.**
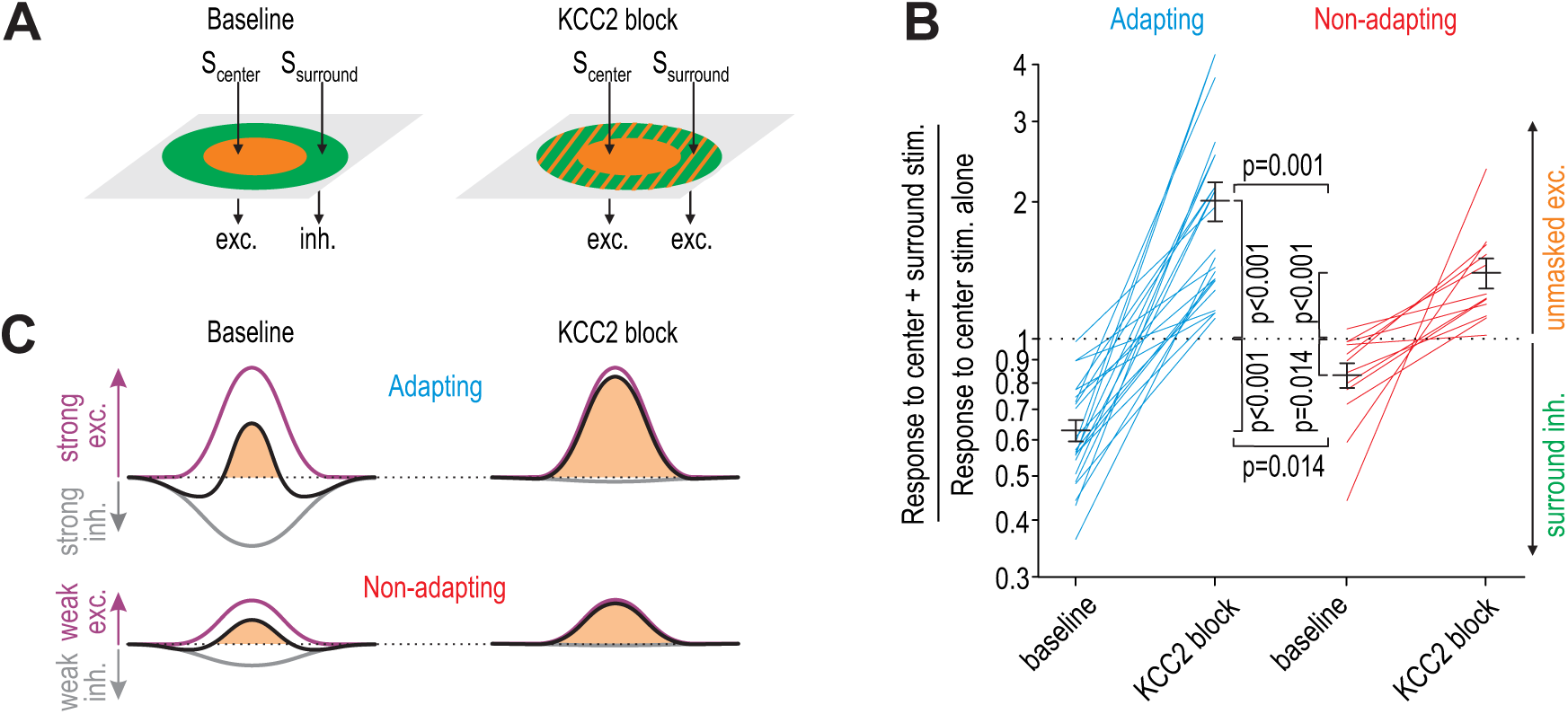
KCC2 blockade unmasks significantly more excitation in adapting units than in non-adapting units. **(A)** Stimulation of the RF center (*S*_center_) evokes net excitation whereas stimulation of the RF surround (*S*_surround_) normally evokes net inhibition but evokes excitation after KCC2 blockade. **(B)** Responses to brush co-stimulation, *S*_center_ + *S*_surround_, expressed as a ratio of the response to *S*_center_ alone. A ratio <1 means that *S*_surround_ evoked inhibition whereas a ratio >1 means that *S*_surround_ contributed to excitation. KCC2 blockade had a significant effect on the ratio (*F*_1,33_ = 86.33, *p* < 0.001; two-way repeated measures ANOVA) but that effect differed between cell types (interaction, *F*_1,33_= 11.30, *p* = 0.002). Specifically, co-stimulation had a significantly stronger inhibitory effect under baseline conditions (*Q*_33_ = 3.56, *p* = 0.014; SNK test) and a significantly stronger excitatory effect after disinhibition (*Q*_33_ = 3.78, *p* = 0.010) in adapting units compared with non-adapting units. **(C)** Cartoon explains data in panel B: adapting units rely on strong inhibition to counterbalance strong touch-evoked excitation whereas non-adapting units require less inhibition to counterbalance weaker excitation. Because of this differential balance of excitation and inhibition, adapting units are disproportionately affected by disinhibition.which was entirely reversed by ACTZ. Results of SNK tests are reported on graphs.

## DISCUSSION

Chloride dysregulation has emerged as an important mechanism by which synaptic inhibition can be compromised (Doyon et al., 2016). It is now well established in dorsal horn neurons that intracellular chloride levels become elevated after nerve injury, that disrupting chloride regulation (e.g. by KCC2 blockade) produces mechanical allodynia, and that restoring chloride regulation reverses the mechanical allodynia caused by nerve injury (for review, see Prescott, 2015). Recent work has verified that these findings apply equally to male and female rodents despite sex-differences in neuroimmune signaling (Mapplebeck et al., in press). Other studies has revealed the diversity of cells types in the dorsal horn and how they connect to form circuits (for reviews, see Peirs and Seal, 2016; Todd, 2017). But many unknowns remain. For instance, without knowing which cell types experience chloride dysregulation and how their spiking is affected, one cannot predict how chloride dysregulation affects circuit function or, in turn, how altered circuit function produces allodynia. Though informative, manipulations targeting a specific cell type fail to address how changes occurring concurrently in different cell types interact at the circuit level. Moreover, the concept of gating, though invaluable in guiding pain research for over 50 years (Melzack and Wall, 1965), is too simplistic to account for complex sensory changes. More detailed investigation of phenomena like spatial and temporal summation is imperative (e.g. Fitzgerald and Jennings, 1999), hence the impetus for the current study and its focus on RF structure.

Our results show that dorsal horn neuron RFs are dramatically enlarged by disinhibition, so much so that after KCC2 blockade, neurons receive more excitatory input from the RF surround than they receive from the RF center (Fig. 1). Though not an initial goal of our study, we were able to separate the majority (∼80%) of the units recorded *in vivo* into two groups that were readily distinguished by their spiking response to tactile stimulation, though several additional features (spike waveform, spontaneous spiking, etc.) also differed between the groups. Pharmacological testing (Fig. 3) as well as the concordance between *in vitro* and *in vivo* testing (Fig. 6E) argue that adapting and non-adapting units correspond to excitatory and inhibitory neurons, respectively. Both cell types experience chloride dysregulation under pathological conditions, modeled here with BDNF (Fig. 7), but certain effects manifest differently.

### Effects of disinhibition on spontaneous spiking

KCC2 blockade (as well as BDNF and SST) increased spontaneous spiking selectively in inhibitory neurons (Figs. 3, 4, and 7). This was reproduced *in vitro* using dynamic clamp, where virtual disinhibition increased the firing rate gain of both cell types but increased spontaneous spiking (reflected in the *y*-intercept) selectively in inhibitory neurons (Fig. 6). Differences in intrinsic cellular properties likely explain the differential effects. First, because of voltage-gated currents, subthreshold depolarization is actively amplified and drives repetitive spiking in tonic-spiking inhibitory neurons but is actively attenuated and fails to drive repetitive spiking in single-spiking excitatory neurons (Prescott and De Koninck, 2005; Ratté et al., 2015). Second, inhibitory neurons are subject to strong tonic inhibition (Gradwell et al., 2017; Takazawa and MacDermott, 2010a). Glycine receptor blockade did not increase spontaneous spiking in inhibitory neurons (Fig. S4) but a strong chloride conductance would make the resting membrane potential of inhibitory neurons very sensitive to KCC2 blockade. Alone or together, these factors render inhibitory neurons prone to spontaneous spiking when chloride is dysregulated.

Excitatory neurons did not spike spontaneously under normal conditions, and so the absence of (increased) spontaneous spiking after KCC2 blockade might be ascribed to a floor effect. However, excitatory neurons responded vigorously to weak tactile stimulation, which means that they do not operate far from spike threshold, contrary to what a floor effect would imply. Moreover, if non-adapting units are indeed inhibitory neurons, the continued lack of spontaneous firing in adapting units after KCC2 blockade argues that GABA/glycinergic transmission does not become paradoxically excitatory, which is consistent with past computational work (Doyon et al., 2011; Prescott et al., 2006) and with past experiments showing that only a small minority of neurons ever exhibit GABA-evoked spikes (Coull et al., 2003).

### Effects of disinhibition on evoked spiking

The effect of chloride dysregulation on evoked spiking is more difficult to interpret because of the many additional factors involved, including the type of primary afferent input (Fig. 2E) and the relative strength of excitation and inhibition triggered by that input, which depends on the circuit. Intrinsic neuronal excitability, reflected in the spiking response to somatic current injection, has also been shown to influence the response to cutaneous stimulation or an *in vitro* facsimile thereof (Graham et al., 2007). At the circuit level, E-I balance depends on where the stimulus occurs relative to the neuron’s RF, which changes over time as a dynamic stimulus moves across an RF. The spatiotemporal characteristics of natural tactile stimuli are not readily reproduced with electrical or optogenetic stimulation. By stimulating different parts of the RF alone or in combination (Fig. 8), our results demonstrate that spatial summation is altered by chloride dysregulation. The consequences of this for sensory processing will depend on the spatial extent of the stimulus on the skin.

A stimulus that extends beyond the RF center and into the RF surround normally engages surround inhibition, causing fewer spikes than if the stimulus was restricted to the RF center. This runs counter to intuition that a broader stimulus will elicit a bigger response, but it is entirely consistent with experiments in the visual system showing that broader stimuli evoke greater inhibition (Haider et al., 2010; Vinje and Gallant, 2002). However, when disinhibition occurs, not only is the surround inhibition lost, so too is subliminal excitatory input unmasked. This means that a stimulus extending into the RF surround will engage more total excitation (and drive stronger spiking) than a stimulus limited to the RF center. Furthermore, beyond one neuron being more strongly activated, more neurons will be activated as their expanded RFs encroach on the area of stimulation. For a projection neuron receiving convergent input from excitatory interneurons, these effects of disinhibition compound – more input from each excitatory interneuron and input from more excitatory interneurons – plus reduced inhibition to the projection neuron itself.

Dynamic allodynia provoked by light brushing of the skin is a far greater clinical problem than the allodynia provoked by punctate stimuli or pressure (Hansson, 2003). In individuals with neuropathic pain, dynamic allodynia is most commonly provoked by clothes moving against the skin, which is precisely the sort of stimulus for which spatial summation is important. That said, preclinical testing of mechanosensitivity relies almost exclusively on von Frey stimulation (Mogil, 2009). Moreover, different forms of mechanical allodynia subserved by different primary afferents (Koltzenburg et al., 1992; Ochoa and Yarnitsky, 1993) and differences in central processing have also been uncovered. Cheng et al. (2017) recently found that dynamic allodynia required a set of neurons defined by the co-expression of VGLUT3 and Lbx1, whereas hypersensitivity to von Frey stimulation persisted after those neurons were ablated. Having tested with both brush and von Frey stimuli, our results show that disinhibition caused by chloride dysregulation affects responses to both types of stimuli, which is consistent with most (if not all) neurons being under inhibitory control. Additional work, including behavioral testing, is required to assess how changes in spatial summation due to chloride dysregulation contributes to allodynia. Human psychophysical testing that extends beyond the standard battery of quantitative sensory tests (Sommer, 2016) would also be very informative.

### Effects of disinhibition on E-I balance

By co-stimulating the RF center and surround and comparing the evoked response with responses to stimulation of the RF center alone (Fig. 8), our results also revealed that the E-I balance differed between putative excitatory and inhibitory neurons. Specifically, we found that excitatory neurons receive strong excitation that is counterbalanced by strong inhibition, whereas inhibitory neurons receive weak excitation that is counterbalanced by weak inhibition. In other words, both cell types appear to receive similar relative levels of excitatory and inhibitory input, but the absolute levels of each input are greater in excitatory neurons. Because excitatory neurons rely on strong inhibition to counterbalance their strong excitatory input, they are more affected than inhibitory neurons by a loss of inhibition. This observation could not have been made through targeted manipulations (e.g. ablation) of a specific cell type, as is now common in many studies (e.g. Cheng et al., 2017). That said, our results are fully consistent with excitatory interneurons playing a critical in allodynia. Consistent with earlier work (Torsney and MacDermott, 2006), our results argue that disinhibition of those excitatory neurons is key.

The disproportionate disinhibition of excitatory neurons is interesting for two reasons. First, the differential effect is due to circuit organization, i.e. the amount of excitatory and inhibitory input received by each cell type. Indeed, we did not observe any difference in chloride regulation between cell types (Fig. 5). Second, sensory processing might not be disrupted if the evoked responses in excitatory and inhibitory neurons were equivalently affected by disinhibition. That said, the disproportionate disinhibition of excitatory neurons is liable to be less evident (or consequential) with punctate stimulation, for which spatial summation is minimal.

In summary, this study has explored the consequences of chloride dysregulation in superficial dorsal horn neurons. Our results show that both excitatory and inhibitory neurons experience chloride dysregulation but respond differently because of differences in their intrinsic cell properties and because of circuit-level differences in E-I balance. Additionally, we demonstrated that dorsal horn neuron RFs have a center-surround organization, and that loss of surround inhibition unmasks substantial excitatory drive, especially in excitatory neurons. Loss of surround inhibition and the unmasking of subliminal excitatory input dramatically alters spatial summation, which may contribute significantly to dynamic allodynia.

## METHODS

All experimental procedures were approved by the Animal Care Committee at the Hospital for Sick Children and were performed in accordance to guidelines from the Canadian Council on Animal Care. Experiments were performed on adult male Sprague Dawley rats obtained from Charles River, Montreal. In separate experiments, we have determined that KCC2 is equally important for chloride regulation in males and female rodents, and that KCC2 downregulation contributes equally to neuropathic pain in both sexes (Mapplebeck et al., in press) despite differences in upstream, neuroimmune signaling (Sorge et al., 2015).

### Animal preparation for in vivo experiments

Under urethane anesthesia (20% in normal saline; 1.2 g/kg i.p.), a laminectomy was performed to expose L4–S1 segments of the spinal cord. The rat (350-450 g) was placed in a stereotaxic frame and its vertebrae clamped above and below the recording site to immobilize the spinal cord. The left hind paw was immobilized in clay with the plantar surface facing upwards for stimulation. Rectal temperature was maintained at 37 °C using a feedback controlled heating pad (TR-200, Fine Science Tools).

### Single Unit Extracellular Recordings

A four-electrode array with a total of 16 recording sites (A4 type, NeuroNexus) was implanted at the L5 spinal level. The array was oriented so that each electrode was at the same mediolateral position. Electrode tips were lowered 328 ± 95 µm (mean ± SD) below the dorsal surface. Since recording sites are spaced at 50 μm intervals up each electrode, we determined on which site a unit was recorded and from this measured the depth of recorded neurons to be <300 µm (see Fig. 3C), which places them within lamina I or II (Watson et al., 2008). Neurons that responded to limb displacement, indicating proprioceptive input, were excluded. The signal was amplified, filtered at 500 Hz – 10 kHz, digitized at 20 kHz with an Omniplex Data Acquisition System (Plexon) and stored with stimulus markers on disk. Single units were isolated using Offline Sorter V3 software (Plexon), and were analyzed with Neuroexplorer 4 (Plexon).

### Drug application in vivo

After the laminectomy, a Vaseline well was built around the spinal cord to target drug delivery to the recording site. In each experiment, 200 µl of saline containing drug was delivered into the well through a small tube (OD: 0.016 inch; Cole-Palmer) inserted intrathecally. To block KCC2, R-(+)-[(2-*n*-butyl-6,7-dichloro-2-cyclopentyl-2,3-dihydro-1-oxo-1H-inden-5-yl)oxy] acetic acid (DIOA; Sigma-Aldrich) and VU0240551 (VU; Tocris Bioscience) were diluted in a stock solution of dimethyl sulphoxide (DMSO) and then diluted in buffered saline for a final concentration of 100 µM for DIOA (Keller et al., 2007) and 50 µM for VU (Lavertu et al., 2014). Results using DIOA and VU were pooled after determining that there was no difference between them, consistent with past work (Lee and Prescott, 2015). 200 µM 3,5-dihydroxyphenylglycine (DHPG; Tocris), 50 µM somatostatin (SST; Abcam), 50 µM strychnine (Abcam) and 70 µg recombinant human brain-derived neurotrophic factor (BDNF; PeproTech) were prepared in saline. Acetazolamide (ACTZ; Sigma-Aldrich) was dissolved in buffered saline with pH 8.2, after which pH was reduced to 7.4 and concentration was adjusted to 10 mM (Asiedu et al., 2009).

### Mechanical stimulation and receptive field mapping

Cutaneous receptive fields (RFs) were identified from the spiking evoked by mechanical stimuli applied by brush, blunt probes, or von Frey filaments to the glabrous skin of the left hind paw. By using weak search stimuli, we targeted neurons receiving low-threshold input and avoided causing sensitization. The RF zone eliciting a response under baseline conditions was defined as the RF center. Testing outside the RF center was conducted prior to any pharmacological manipulations (to demonstrate the absence of suprathreshold input) but the RF surround was only mapped after KCC2 blockade. Each stimulus comprised ten 1 s-long applications of the brush or von Frey filament (2, 4, 6, 8, and 10 g) repeated at 2 s intervals. Each stimulus was applied twice onto different locations within each RF zone for each test condition. For experiments in Figure 8, two brushes were held together and applied concurrently to two different RF zones.

### In vivo data analysis

To quantify the spiking evoked by mechanical stimulation, we measured the mean firing rate during each stimulus, subtracted the spontaneous firing rate measured from the 10-s epoch preceding each stimulus, and then averaged across the two stimuli for each RF zone and test condition. Input-output (*i-o*) plots produced by varying von Frey force were fitted by linear regression to obtain the slope for each neuron. All data are presented as mean ± SEM.

### Slice preparation for in vitro experiments

Spinal slices were prepared as previously described (Prescott and De Koninck, 2002). Briefly, the rat (100-200 g) was anesthetized with 4% isoflorane and perfused intracardially with ice-cold oxygenated (95% O_2_ and 5% CO_2_) sucrose-substituted artificial cerebrospinal fluid (ACSF) containing (in mM): 252 sucrose, 2.5 KCl, 2 CaCl_2_, 2 MgCl_2_, 10 glucose, 26 NaHCO_3_, 1.25 NaH_2_PO_4_ and 5 kynurenic acid; pH 7.35. The rat was decapitated, the spinal cord was removed by hydraulic extrusion, and 300–400 μm-thick sections were cut from the lumbar enlargement in the parasagittal plane. Slices were kept in normal oxygenated ACSF (126 mM NaCl instead of sucrose and without kynurenic acid) at room temperature until recording.

### Whole-cell patch clamp recordings

Slices were transferred to a recording chamber constantly perfused at ∼2 ml/min with oxygenated (95% O_2_ and 5% CO_2_) ACSF. Visually identified neurons in lamina I and II were patched in the whole cell configuration with >70% series resistance compensation using an Axopatch 200B amplifier (Molecular Devices). Patch pipettes were pulled from borosilicate glass capillaries (WPI) and were filled with one of two intracellular solutions. For dynamic clamp experiments, the solution comprised (in mM): 125 KMeSO_4_, 5 KCl, 10 HEPES, 2 MgCl_2_, 4 ATP, 0.4 GTP as well as 0.1% Lucifer Yellow; pH was adjusted to 7.2 with KOH. For chloride extrusion measurements, KCl was increased to 25 mM and KMeSO_4_ was reduced to 115 mM. Recordings were at room temperature (22–26 °C). Traces were low-pass filtered at 2 KHz, digitized at 20 KHz using a CED 1401 computer interface (Cambridge Electronic Design), and analyzed offline. Membrane potential (after correction for the liquid junction potential) was adjusted to −65 mV through tonic current injection. Based on responses to series of 500 ms-long current steps, spiking patterns were classified as previously described (Prescott and De Koninck, 2002).

### Chloride extrusion measurements

Spiking pattern was determined in current clamp mode immediately after rupturing the membrane. Thereafter, 1 µM tetrodotoxin (Alomone Labs) and 1 µM CGP55845 (Abcam) were bath applied to block sodium and GABA_B_ channels, respectively. After allowing at least 15 min for the chloride from the pipette solution to dialyze, the cell was voltage clamped at different holding potentials while short (5-10 ms-long) puffs of GABA (1 mM prepared in HEPES-buffered pipette solution) were applied ∼5 µm from the soma using a puff pipette (5 MΩ resistance) attached to a picospritzer (Toohey Company, Fairfield, NJ). Experiments were repeated after bath application of 15 µM VU to block KCC2.

### Virtual synaptic input by dynamic clamp

To simulate irregular synaptic input, noisy excitatory and inhibitory conductances (*g*_exc_ and *g*_inh_) were constructed from separate Ornstein-Uhlenbeck processes (Prescott and De Koninck, 2009; Uhlenbeck and Ornstein, 1930) such that *g* fluctuates randomly while returning to its average value *g*_0_ with a time constant *τ* according to

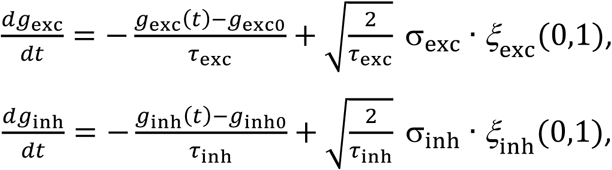

where ξ(0,1) is a random number with 0 mean and unit variance and 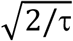is a scaling factor so that σ specifies the standard deviation of the conductance fluctuations. Based on synaptic decay kinetics, *τ*_exc_ = 3 ms and *τ*_inh_ = 10 ms give appropriate autocorrelation structure. Conductances were rectified to disallow negative values. The voltage *V* is recorded from the neuron at each time step, from which the driving forces are calculated and multiplied by the time-varying conductances, and the resultant currents summed according to

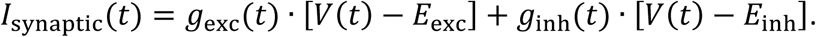

*I*_synaptic_ was applied to the recorded neuron using the dynamic clamp capability of Signal 5 software (Cambridge Electronic Design). Input-output (*i-o*) curves were plotted by varying *g*_exc0_ while co-varying *g*_inh0_ according to a fixed ratio of inhibition to excitation, where α = *g*_inh0_/*g*_exc0_ = 0.5 or 2 to simulate stimulation in the RF center or surround, respectively. *E*_inh_ was set to −70 mV or −45 mV to simulate normal and disinhibited conditions, respectively. *E*_exc_ = 0 mV. Background synaptic activity was blocked with 10 µM bicuculline and 10 µM CNQX (Abcam).

### Statistical Analysis

All statistical analysis was conducted in SigmaPlot 11. An appropriate non-parametric test was applied when the normality (Shapiro-Wilk) test failed. Paired tests were used in all cases where responses from the same cell were measured before and after a manipulation. Unpaired tests were used for between-group comparisons. All tests were two-sided. Once a recorded neuron was classified based on its spiking pattern (see Results) and was included or excluded on the basis of that pattern, no data were excluded for any other reason. Sample sizes are evident on the figures, which show data from individual cells, and from the degrees of freedom reported with statistical tests. Sample sizes were not computed because we did not have any prior knowledge of the expected effect size or variance; instead, experiments were continued until we collected >*n* cells of each type for each experiment, where *n* = 5 for *in vitro* experiments and *n* = 10 for *in vivo* experiments (because variability is typically greater *in vivo*). The only exception is in Figure 2D, where data collected for analysis in Figure 1 were re-analyzed after classification described in Figure 2B. The total number or type of cells recorded from each rat is unpredictable, but cells comprising each data set come from ≥5 rats in all cases.

## ACKNOWLEDGEMENTS

We thank Y De Koninck and MW Salter for their feedback on an early version of this manuscript. This work was supported by a Project Grant from the Canadian Institutes of Health Research (PJT-153161).

---

**Figure S4.**
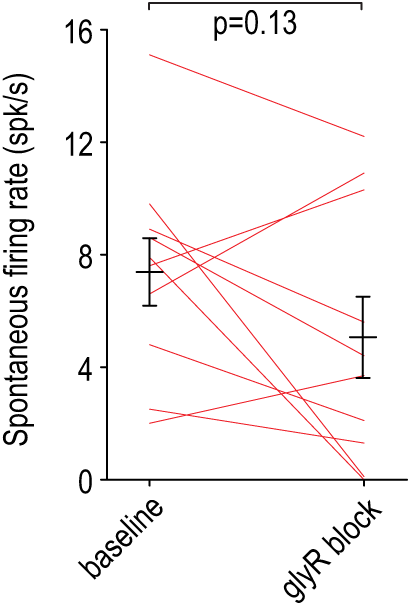
Effect of glycine receptor blockade on spontaneous spiking in non-adapting units. Intrathecal strychnine (50 µM) had no significant effect on spontaneous spiking (*T*_9_ = 1.65, *p* = 0.13; paired *t*-test).

**Figure S6.**
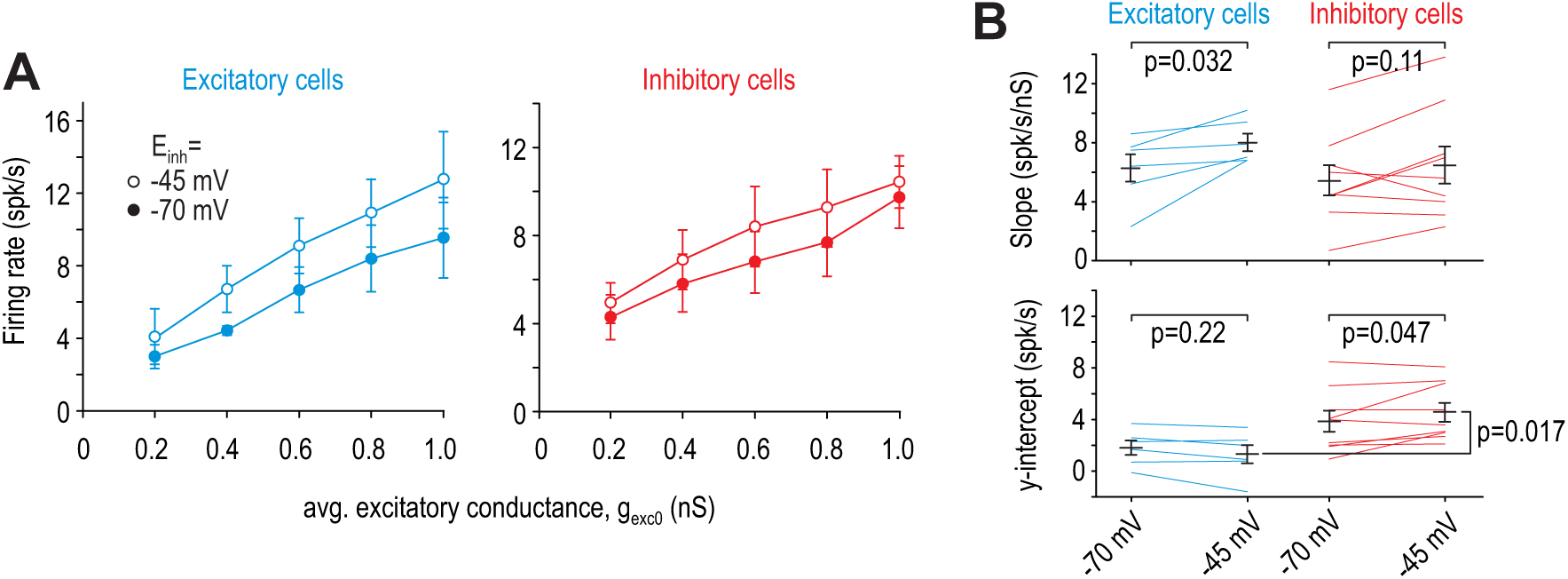
Effects of virtual disinhibition for α = 0.5 (virtual stimulation in RF centre). **(A)** Average *i-o* curves (±SEM) for excitatory neurons (blue) and inhibitory neurons (red). **(B)** *E*_inh_ had a significant effect on slope (*F*_1,13_= 8.82, *p* = 0.011, two-way repeated measures ANOVA) but only in excitatory cells. Results of SNK tests are reported on graphs. The effect of *E*_inh_ on the *y*-intercept differed significantly between excitatory and inhibitory cells (interaction, *F*_1,13_ = 5.67, *p* = 0.033), with only inhibitory cells experiencing a significant change. Like for virtual stimulation in the RF surround (Fig. 6), the *y*-intercept differed significantly (*p* = 0.017; SNK test) between excitatory and inhibitory cells for *E*_inh_ = −45 mV.

## REFERENCES

Abraira, V.E., Kuehn, E.D., Chirila, A.M., Springel, M.W., Toliver, A.A., Zimmerman, A.L., Orefice, L.L., Boyle, K.A., Bai, L., Song, B.J., et al. (2017). The Cellular and Synaptic Architecture of the Mechanosensory Dorsal Horn. Cell 168, 295-310.e219.

Asiedu, M., Ossipov, M.H., Kaila, K., and Price, T.J. (2009). Acetazolamide and midazolam act synergistically to inhibit neuropathic pain. Pain 148, 302–308.

Baba, H., Ji, R.R., Kohno, T., Moore, K.A., Ataka, T., Wakai, A., Okamoto, M., and Woolf, C.J. (2003). Removal of GABAergic inhibition facilitates polysynaptic A fiber-mediated excitatory transmission to the superficial spinal dorsal horn. Mol Cell Neurosci 24, 818–830.

Balachandar, A., and Prescott, S.A. (2018). Origin of heterogeneous spiking patterns from continuously distributed ion channel densities: a computational study in spinal dorsal horn neurons. J Physiol 596, 1681–1697.

Behbehani, M.M., and Dollberg-Stolik, O. (1994). Partial sciatic nerve ligation results in an enlargement of the receptive field and enhancement of the response of dorsal horn neurons to noxious stimulation by an adenosine agonist. Pain 58, 421–428.

Cheng, L., Duan, B., Huang, T., Zhang, Y., Chen, Y., Britz, O., Garcia-Campmany, L., Ren, X., Vong, L., Lowell, B.B., et al. (2017). Identification of spinal circuits involved in touch-evoked dynamic mechanical pain. Nat Neurosci 20, 804–814.

Cook, A.J., Woolf, C.J., Wall, P.D., and McMahon, S.B. (1987). Dynamic receptive field plasticity in rat spinal cord dorsal horn following C-primary afferent input. Nature 325, 151–153.

Coull, J.A., Boudreau, D., Bachand, K., Prescott, S.A., Nault, F., Sik, A., De Koninck, P., and De Koninck, Y. (2003). Trans-synaptic shift in anion gradient in spinal lamina I neurons as a mechanism of neuropathic pain. Nature 424, 938–942.

Coull, J.A.M., Beggs, S., Boudreau, D., Boivin, D., Tsuda, M., Inoue, K., Gravel, C., Salter, M.W., and De Koninck, Y. (2005). BDNF from microglia causes the shift in neuronal anion gradient underlying neuropathic pain. Nature 438, 1017–1021.

Cumberbatch, M.J., Carlson, E., Wyatt, A., Boyce, S., Hill, R.G., and Rupniak, N.M. (1998). Reversal of behavioural and electrophysiological correlates of experimental peripheral neuropathy by the NK1 receptor antagonist GR205171 in rats. Neuropharmacology 37, 1535–1543.

Devor, M., and Wall, P.D. (1981). Effect of peripheral nerve injury on receptive fields of cells in the cat spinal cord. J Comp Neurol 199, 277–291.

Doyon, N., Prescott, S.A., Castonguay, A., Godin, A.G., Kroger, H., and De Koninck, Y. (2011). Efficacy of synaptic inhibition depends on multiple dynamically interacting mechanisms implicated in chloride homeostasis. PLoS Comput Biol 7, e1002149.

Doyon, N., Vinay, L., Prescott, S.A., and De Koninck, Y. (2016). Chloride Regulation: A Dynamic Equilibrium Crucial for Synaptic Inhibition. Neuron 89, 1157–1172.

Duan, B., Cheng, L., Bourane, S., Britz, O., Padilla, C., Garcia-Campmany, L., Krashes, M., Knowlton, W., Velasquez, T., Ren, X., et al. (2014). Identification of spinal circuits transmitting and gating mechanical pain. Cell 159, 1417–1432.

Dubuisson, D., Fitzgerald, M., and Wall, P.D. (1979). Ameboid receptive fields of cells in laminae 1, 2 and 3. Brain Res 177, 376–378.

Fitzgerald, M., and Jennings, E. (1999). The postnatal development of spinal sensory processing. Proc Natl Acad Sci USA 96, 7719–7722.

Gagnon, M., Bergeron, M.J., Lavertu, G., Castonguay, A., Tripathy, S., Bonin, R.P., Perez-Sanchez, J., Boudreau, D., Wang, B., Dumas, L., et al. (2013). Chloride extrusion enhancers as novel therapeutics for neurological diseases. Nat Med 19, 1524–1528.

Gradwell, M.A., Boyle, K.A., Callister, R.J., Hughes, D.I., and Graham, B.A. (2017). Heteromeric alpha/beta glycine receptors regulate excitability in parvalbumin-expressing dorsal horn neurons through phasic and tonic glycinergic inhibition. J Physiol 595, 7185–7202.

Graham, B.A., Brichta, A.M., and Callister, R.J. (2007). Pinch-current injection defines two discharge profiles in mouse superficial dorsal horn neurones, in vitro. J Physiol 578, 787–798.

Haider, B., Krause, M.R., Duque, A., Yu, Y., Touryan, J., Mazer, J.A., and McCormick, D.A. (2010). Synaptic and network mechanisms of sparse and reliable visual cortical activity during nonclassical receptive field stimulation. Neuron 65, 107–121.

Hansson, P. (2003). Difficulties in stratifying neuropathic pain by mechanisms. EurJ Pain 7, 353–357.

Hillman, P., and Wall, P.D. (1969). Inhibitory and excitatory factors influencing the receptive fields of lamina 5 spinal cord cells. Exp Brain Res 9, 284–306.

Hu, H.J., Carrasquillo, Y., Karim, F., Jung, W.E., Nerbonne, J.M., Schwarz, T.L., and Gereau, R.W. (2006). The kv4.2 potassium channel subunit is required for pain plasticity. Neuron 50, 89–100.

Hu, H.J., and Gereau, R.W. (2011). Metabotropic glutamate receptor 5 regulates excitability and Kv4.2-containing K channels primarily in excitatory neurons of the spinal dorsal horn. J Neurophysiol 105, 3010–3021.

Hughes, D.I., Sikander, S., Kinnon, C.M., Boyle, K.A., Watanabe, M., Callister, R.J., and Graham, B.A. (2012). Morphological, neurochemical and electrophysiological features of parvalbumin-expressing cells: a likely source of axo-axonic inputs in the mouse spinal dorsal horn. J Physiol 590, 3927–3951.

Isaacson, J.S., and Scanziani, M. (2011). How inhibition shapes cortical activity. Neuron 72, 231–243.

Johansson, R.S., and Vallbo, A.B. (1979). Tactile sensibility in the human hand: relative and absolute densities of four types of mechanoreceptive units in glabrous skin. J Physiol 286, 283–300.

Keller, A.F., Beggs, S., Salter, M.W., and De Koninck, Y. (2007). Transformation of the output of spinal lamina I neurons after nerve injury and microglia stimulation underlying neuropathic pain. Mol Pain 3, 27.

Koltzenburg, M., Lundberg, L.E., and Torebjork, H.E. (1992). Dynamic and static components of mechanical hyperalgesia in human hairy skin. Pain 51, 207–219.

Kuffler, S.W. (1953). Discharge patterns and functional organization of mammalian retina. J Neurophysiol 16, 37–68.

Laird, J.M., and Cervero, F. (1989). A comparative study of the changes in receptive-field properties of multireceptive and nocireceptive rat dorsal horn neurons following noxious mechanical stimulation. J Neurophysiol 62, 854–863.

Lavertu, G., Cote, S.L., and De Koninck, Y. (2014). Enhancing K-Cl co-transport restores normal spinothalamic sensory coding in a neuropathic pain model. Brain 137, 724–738.

Lee, K.Y., and Prescott, S.A. (2015). Chloride dysregulation and inhibitory receptor blockade yield equivalent disinhibition of spinal neurons yet are differentially reversed by carbonic anhydrase blockade. Pain 156, 2431–2437.

Lu, Y., Dong, H., Gao, Y., Gong, Y., Ren, Y., Gu, N., Zhou, S., Xia, N., Sun, Y.Y., Ji, R.R., and Xiong, L. (2013). A feed-forward spinal cord glycinergic neural circuit gates mechanical allodynia. J Clin Invest 123, 4050–4062.

Malmberg, A.B., Chen, C., Tonegawa, S., and Basbaum, A.I. (1997). Preserved acute pain and reduced neuropathic pain in mice lacking PKCgamma. Science 278, 279–283.

Mapplebeck, J.C., Lorenzo, L.E., Lee, K.Y., Gauthier, C., Mulley, M.M., De Koninck, Y., Prescott, S.A., and Salter, M.W. (in press). Chloride dysregulation through downregulation of KCC2 mediates neuropathic pain in both sexes. Cell Rep.

McMahon, S.B., and Wall, P.D. (1984). Receptive fields of rat lamina 1 projection cells move to incorporate a nearby region of injury. Pain 19, 235–247.

Melzack, R., and Wall, P.D. (1965). Pain mechanisms: a new theory. Science 150, 971–979.

Miraucourt, L.S., Dallel, R., and Voisin, D.L. (2007). Glycine inhibitory dysfunction turns touch into pain through PKCgamma interneurons. PLoSONE 2, e1116.

Miraucourt, L.S., Moisset, X., Dallel, R., and Voisin, D.L. (2009). Glycine inhibitory dysfunction induces a selectively dynamic, morphine-resistant, and neurokinin 1 receptor-independent mechanical allodynia. J Neurosci 29, 2519–2527.

Mogil, J.S. (2009). Animal models of pain: progress and challenges. NatRevNeurosci 10, 283–294.

Ochoa, J.L., and Yarnitsky, D. (1993). Mechanical hyperalgesias in neuropathic pain patients: dynamic and static subtypes. Ann Neurol 33, 465–472.

Peirs, C., and Seal, R.P. (2016). Neural circuits for pain: Recent advances and current views. Science 354, 578–584.

Pleger, B., Ragert, P., Schwenkreis, P., Forster, A.F., Wilimzig, C., Dinse, H., Nicolas, V., Maier, C., and Tegenthoff, M. (2006). Patterns of cortical reorganization parallel impaired tactile discrimination and pain intensity in complex regional pain syndrome. Neuroimage 32, 503–510.

Polgar, E., Hughes, D.I., Riddell, J.S., Maxwell, D.J., Puskar, Z., and Todd, A.J. (2003). Selective loss of spinal GABAergic or glycinergic neurons is not necessary for development of thermal hyperalgesia in the chronic constriction injury model of neuropathic pain. Pain 104, 229–239.

Prescott, S.A. (2015). Synaptic inhibition and disinhibition in the spinal dorsal horn. In Molecular Biology of Pain, Progress in Molecular and Translational Science, T.J. Price, and G. Dussor, eds. (Elsevier).

Prescott, S.A., and De Koninck, Y. (2002). Four cell types with distinctive membrane properties and morphologies in lamina I of the spinal dorsal horn of the adult rat. JPhysiol 539, 817–836.

Prescott, S.A., and De Koninck, Y. (2003). Gain control of firing rate by shunting inhibition: roles of synaptic noise and dendritic saturation. Proc Natl Acad Sci USA 100, 2076–2081.

Prescott, S.A., and De Koninck, Y. (2005). Integration time in a subset of spinal lamina I neurons is lengthened by sodium and calcium currents acting synergistically to prolong subthreshold depolarization. J Neurosci 25, 4743–4754.

Prescott, S.A., and De Koninck, Y. (2009). Impact of background synaptic activity on neuronal response properties revealed by stepwise replication of in vivo -like conditions in vitro In The Dynamic Clamp: From Principles to Applications, A. Destexhe, and T. Bal, eds. (Springer), pp. 89–114.

Prescott, S.A., Ma, Q., and De Koninck, Y. (2014). Normal and abnormal coding of somatosensory stimuli causing pain. Nat Neurosci 17, 183–191.

Prescott, S.A., and Ratté, S. (2012). Pain processing by spinal microcircuits: afferent combinatorics. Curr Opin Neurobiol 22, 631–639.

Prescott, S.A., Sejnowski, T.J., and De Koninck, Y. (2006). Reduction of anion reversal potential subverts the inhibitory control of firing rate in spinal lamina I neurons: towards a biophysical basis for neuropathic pain. Mol Pain 2, 32.

Price, T.J., Cervero, F., Gold, M.S., Hammond, D.L., and Prescott, S.A. (2009). Chloride regulation in the pain pathway. Brain Res Rev 60, 149–170.

Ratté, S., Lankarany, M., Rho, Y.A., Patterson, A., and Prescott, S.A. (2015). Subthreshold membrane currents confer distinct tuning properties that enable neurons to encode the integral or derivative of their input. Front Cell Neurosci 8, 452.

Silver, R.A. (2010). Neuronal arithmetic. Nat Rev Neurosci 11, 474–489.

Sivilotti, L., and Woolf, C.J. (1994). The contribution of GABA(A) and glycine receptors to central sensitization: disinhibition and touch-evoked allodynia in the spinal cord. J Neurophysiol 72, 169–179.

Sommer, C. (2016). Exploring pain pathophysiology in patients. Science 354, 588–592.

Sorge, R.E., Mapplebeck, J.C., Rosen, S., Beggs, S., Taves, S., Alexander, J.K., Martin, L.J., Austin, J.S., Sotocinal, S.G., Chen, D., et al. (2015). Different immune cells mediate mechanical pain hypersensitivity in male and female mice. Nat Neurosci 18, 1081–1083.

Sorkin, L.S., and Puig, S. (1996). Neuronal model of tactile allodynia produced by spinal strychnine: effects of excitatory amino acid receptor antagonists and a mu-opiate receptor agonist. Pain 68, 283–292.

Sorkin, L.S., Puig, S., and Jones, D.L. (1998). Spinal bicuculline produces hypersensitivity of dorsal horn neurons: effects of excitatory amino acid antagonists. Pain 77, 181–190.

Spike, R.C., Puskar, Z., Andrew, D., and Todd, A.J. (2003). A quantitative and morphological study of projection neurons in lamina I of the rat lumbar spinal cord. Eur J Neurosci 18, 2433–2448.

Suzuki, R., Kontinen, V.K., Matthews, E., Williams, E., and Dickenson, A.H. (2000). Enlargement of the receptive field size to low intensity mechanical stimulation in the rat spinal nerve ligation model of neuropathy. ExpNeurol 163, 408–413.

Tabo, E., Jinks, S.L., Eisele, J.H., Jr., and Carstens, E. (1999). Behavioral manifestations of neuropathic pain and mechanical allodynia, and changes in spinal dorsal horn neurons, following L4-L6 dorsal root constriction in rats. Pain 80, 503–520.

Takazawa, T., and MacDermott, A.B. (2010a). Glycinergic and GABAergic tonic inhibition fine tune inhibitory control in regionally distinct subpopulations of dorsal horn neurons. J Physiol 588, 2571–2587.

Takazawa, T., and MacDermott, A.B. (2010b). Synaptic pathways and inhibitory gates in the spinal cord dorsal horn. Ann N Y Acad Sci 1198, 153–158.

Todd, A.J. (2010). Neuronal circuitry for pain processing in the dorsal horn. Nat Rev Neurosci 11, 823–836.

Todd, A.J. (2017). Identifying functional populations among the interneurons in laminae I-III of the spinal dorsal horn. Mol Pain 13, 1744806917693003.

Todd, A.J., Spike, R.C., and Polg r, E. (1998). A quantitative study of neurons which express neurokinin-1 or somatostatin sst2a receptor in rat spinal dorsal horn. Neuroscience 85, 459–473.

Torsney, C., and MacDermott, A.B. (2006). Disinhibition opens the gate to pathological pain signaling in superficial neurokinin 1 receptor-expressing neurons in rat spinal cord. J Neurosci 26, 1833–1843.

Uhlenbeck, G.E., and Ornstein, L.S. (1930). On the theory of Brownian motion. PhysRev 36, 823–841.

van Hecke, O., Austin, S.K., Khan, R.A., Smith, B.H., and Torrance, N. (2014). Neuropathic pain in the general population: a systematic review of epidemiological studies. Pain 155, 654–662.

Vinje, W.E., and Gallant, J.L. (2002). Natural stimulation of the nonclassical receptive field increases information transmission efficiency in V1. J Neurosci 22, 2904–2915.

Wang, X., Zhang, J., Eberhart, D., Urban, R., Meda, K., Solorzano, C., Yamanaka, H., Rice, D., and Basbaum, A.I. (2013). Excitatory superficial dorsal horn interneurons are functionally heterogeneous and required for the full behavioral expression of pain and itch. Neuron 78, 312–324.

Watson, C., Paxinos, G., and Kayalioglu, G. (2008). The Spinal Cord: A Christopher and Dana Reeve Foundation Text and Atlas (New York: Elsevier).

Woolf, C.J. (2010). Overcoming obstacles to developing new analgesics. Nat Med 16, 1241–1247.

Yaksh, T.L. (1989). Behavioral and autonomic correlates of the tactile evoked allodynia produced by spinal glycine inhibition: effects of modulatory receptor systems and excitatory amino acid antagonists. Pain 37, 111–123.

Yasaka, T., Tiong, S.Y., Hughes, D.I., Riddell, J.S., and Todd, A.J. (2010). Populations of inhibitory and excitatory interneurons in lamina II of the adult rat spinal dorsal horn revealed by a combined electrophysiological and anatomical approach. Pain 151, 475–488.

Zeilhofer, H.U., Wildner, H., and Yevenes, G.E. (2012). Fast synaptic inhibition in spinal sensory processing and pain control. Physiol Rev 92, 193–235.

Zheng, J., Lu, Y., and Perl, E.R. (2010). Inhibitory neurones of the spinal substantia gelatinosa mediate interaction of signals from primary afferents. J Physiol 588, 2065–2075.

